# Small proteins modulate ion channel-like ACD6 to regulate immunity in *Arabidopsis thaliana*

**DOI:** 10.1101/2021.01.25.428077

**Authors:** Junbin Chen, Lei Li, Jong Hum Kim, Benjamin Neuhäuser, Mingyu Wang, Michael Thelen, Richard Hilleary, Yuan Chi, Luyang Wei, Kavita Venkataramani, Moises Exposito-Alonso, Chang Liu, Jakob Keck, A. Cristina Barragan, Rebecca Schwab, Ulrich Lutz, Zhen-Ming Pei, Sheng-Yang He, Uwe Ludewig, Detlef Weigel, Wangsheng Zhu

**Affiliations:** China Key Laboratory of Key Laboratory of Surveillance and Management for Plant Quarantine Pests, MARA, and College of Plant Protection, China Agricultural University, 100193, Beijing, China; Department of Molecular Biology, Max Planck Institute for Biology Tübingen, 72076 Tübingen, Germany; Department of Biology, Duke University, Durham, NC 27708, USA; Nutritional Crop Physiology, University of Hohenheim, 70599 Stuttgart, Germany; Center for Plant Molecular Biology (ZMBP), University of Tübingen, 72076 Tübingen, Germany; Institute of Biology, University of Hohenheim, 70599 Stuttgart, Germany; Howard Hughes Medical Institute, Duke University, Durham, NC 27708, USA; Institute for Genetics and Developmental Biology, Chinese Academy of Sciences, 100101 Beijing, China; Institute for Physical and Theoretical Chemistry, University Tübingen, 72076 Tübingen, Germany; Department of Biology, University of Massachusetts Boston, Boston, MA 02125, USA; Department of Plant Biology, Carnegie Institution for Science, Stanford, CA 94305, USA; The Sainsbury Laboratory, Norwich Research Park, Norwich NR4 7UH, UK

**Author notes:** Correspondence (W.Z.); (D.W.). These authors contributed equally (Junbin Chen, Lei Li, Jong Hum Kim, Benjamin Neuhäuser).

## Abstract

ACCELERATED CELL DEATH 6 (ACD6) mediates a trade-off between growth and defense in *Arabidopsis thaliana*. However, the precise biochemical mechanism by which ACD6 and related proteins in plants act remains enigmatic. Here, we identified two loci, *MODULATOR OF HYPERACTIVE ACD6 1* (*MHA1*) and its paralog *MHA1-LIKE* (*MHA1L*), that code for ∼7 kDa proteins that differentially interact with specific ACD6 variants. MHA1L enhances accumulation of an ACD6 complex, thereby increasing activity of the *ACD6* standard allele for regulating plant growth and defenses. ACD6 is a multipass transmembrane protein with intracellular ankyrin repeats that are structurally similar to those found in mammalian ion channels. Several lines of evidence link increased ACD6 activity to enhanced calcium influx, likely mediated by ACD6 itself and with MHA1L as a direct regulator of ACD6.

## INTRODUCTION

While plants need to defend themselves against pathogens, an inherent danger is inappropriate activation of immune responses, which can cause collateral damage to the plant itself. Particularly effective immune alleles enable a plant to respond rapidly to pathogen attack, but such variants might also be potentially deleterious. The study of natural variation in the immune system of *Arabidopsis thaliana* has identified *ACCELERATED CELL DEATH 6* (*ACD6*), a positive regulator of cell death and defense responses, as a nexus for a trade-off between growth and disease resistance in wild populations^1–3^. The natural *ACD6*-Est-1 allele can protect plants against a wide range of unrelated pathogens, but at the same time often exacts a substantial growth penalty in form of reduced stature and cell death in leaves^1^.

The hyperimmunity conditioned by high levels of *ACD6* activity is dependent on salicylic acid (SA), with inactivation of individual SA signaling components partially suppressing the effects of *ACD6* hyperactivity^4, 5^. SA signaling is required for several aspects of plant immunity, including pattern- and effector-triggered immunity (PTI and ETI) as well as systemic acquired resistance (SAR)^6–9^. Both SA biosynthesis and many aspects of SA up- and downstream signaling are well understood, with feedback and feedforward regulation being important mechanisms. *ACD6* stimulates SA accumulation, and SA in turn enhances *ACD6* mRNA expression and affects the subcellular localization of ACD6 protein^10, 11^. A working model for ACD6 is that it is a redundant regulator of SA signaling through a positive SA-dependent feedback loop^10–12^. An *ACD6* homolog, *BDA1*, also functions as a positive regulator of immunity in *A. thaliana*^13^, as do maize and wheat *ACD6* homologs, which have recently been implicated in smut (*ZmACD6*) and leaf-rust (*Lr14a*) resistance, respectively^14, 15^.

*ACD6* encodes a multipass transmembrane protein with nine intracellular ankyrin repeats^10, 11^. ACD6 protein has been found in association with multiple plasma membrane-localized pattern recognition receptors (PRRs) and the PRR co-receptor BRI1-ASSOCIATED RECEPTOR KINASE 1 (BAK1) as well as other receptor like kinases, supporting a direct role for ACD6 in plant immunity^11, 12, 16^. In response to SA signaling, ACD6 oligomers accumulate at the plasma membrane, indicating that plasma membrane localisation of ACD6 is important for its function^11, 12^. *ACD6* interacts genetically also with the TIR-NLR receptor gene *SUPPRESSOR OF NPR1-1, CONSTITUTIVE 1* (*SNC1*), with the *SNC1* weak allele attenuating the atutoimmune phenotype of *ACD6*-Est-1 in some wild strains of *Arabidopsis*^17^. The ACD6 homolog Lr14a in wheat could trigger Lanthanum(III) chloride inhibited water-soaking phenotype in *N.benthamiana* leaf^15^, suggesting role of ACD6 homologs in regulating calcium fluxes. However, the precise biochemical mechanism by which ACD6 and related proteins act remains enigmatic.

There has recently been considerable progress about a series of plant immune proteins that regulate calcium influx during pathogen responses^18^. These include classical ion channels, such as glutamate receptor-like proteins (GLRs)^19^, cyclic nucleotide-gated ion channels (CNGCs)^20^, reduced hyperosmolality-induced [Ca^2+^]_cyt_ increase channels (OSCAs)^21^, as well as non-canonical NLR-formed ion channels^22, 23^. Here, starting from the analysis of natural modifiers of *ACD6* activity, we identify a new gene family encoding small proteins, MODIFIER OF HYPERACTIVE ACD6 1 (MHA1) and its paralog MHA1-LIKE (MHA1L), which affect ACD6 activity in a complex manner. Sequence and structure similarity of ankyrin repeats in ACD6 with those of transient receptor protein (TRP) ion channels from animals and fungi spurred us to investigate whether MHA1 and/or MHA1L effects might be mediated by ion-dependent ACD6 signaling. We find that increased ACD6 activity enhances calcium signaling, most likely because ACD6 itself is an ion channel. Since MHA1 and MHA1L both bind to the ankyrin repeats of ACD6, we propose that MHA1 and MHA1L are ACD6 ligands that regulate ACD6-dependent ion channel activity and signaling.

## RESULTS

### Identification of MODIFIER OF HYPERACTIVE ACD6 (MHA) loci

Many, but not all *A. thaliana* accessions carrying the *ACD6*-Est-1 allele express visible signs of autoimmunity in the absence of pathogen challenge^1^. Using genome-wide association study (GWAS) of accessions with the *ACD6*-Est-1 allele, we identified two unlinked regions, *MODIFIER OF HYPERACTIVE ACD6 1* (*MHA1*) (Chr1: 22,935,037, *p*=10^-12^), and *MHA2* (Chr4: 11,019,243, *p*=10^-8^) **(Figure 1A)**, which together explained over 60% of variation in macroscopic cell death among *ACD6*-Est-1 carriers. To confirm the GWAS results, we made use of the accession Ty-0, which has cell-death suppressing alleles at both *MHA1* and *MHA2* **(Figures 1B and S1A)**. There were no nonsynonymous differences in *ACD6* between Ty-0 and Est-1, and *ACD6* was well expressed **(Figure S1B)**, but Ty-0 had much less SA than Est-1, and the SA-responsive marker gene *PATHOGENESIS-RELATED 1* (*PR1*) was also expressed at much lower levels **(Figures S1B and S1C)**. In support of phenotypic differences mapping outside of *ACD6*, *ACD6*-Ty-0 and *ACD6*-Est-1 similarly triggered necrotic lesions and reduced biomass when introduced as transgenes into the Col-0 reference background **(Figures S1D and S1E)**. Together, these results demonstrate that Ty-0 harbors extragenic suppressors that modulate the activity of *ACD6*-Est-1.

**Figure 1.**
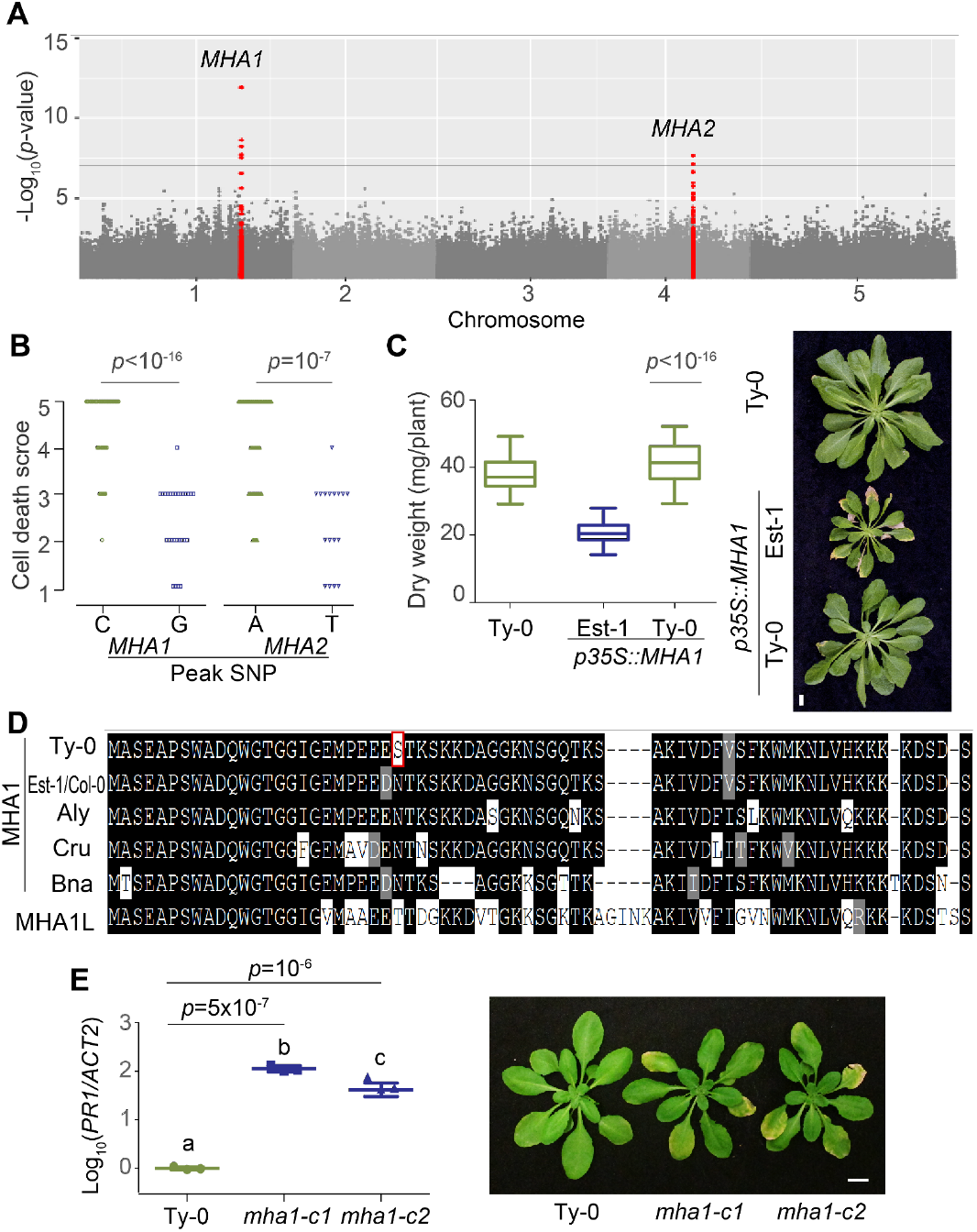
Identification of *MHA1* as a genetic modifier of *ACD6-*Est-1. **(A)** Manhattan plot of GWAS scores for cell death severity. SNPs within 60 kb of the top SNPs in red. The horizontal gray line indicates a nominal *p*=0.05 threshold after Bonferroni correction. **(B)** Cell death severity score of accessions with different *MHA1* and *MHA2* alleles at peak SNP from GWAS analysis shown in (A) (*p*-value from Wilcoxon nonparametric test). **(C)** Left, Dry weight of 4-week-old plants with different *MHA1* alleles. *p*-values from Tukey’s HSD test. right, Five-week-old *p35S::MHA1* lines in Ty-0 background grown in 23°C SD. Top, non-transgenic control. **(D)** MHA1 and MHA1L protein alignments among Brassicaceae (*Aly*, *Arabidopsis lyrata*; *Cru*, *Capsella rubella*; *Bna*, *Brassica napus*). Ty-0-specific polymorphism in MHA1 highlighted in red. **(E)** Left: Accumulation of *PR1* mRNA, as measured by qRT-PCR from three biological replicates each in Ty-0 *mha1* mutants. Right: Plant phenotypes. Scale bar, 1 cm. **See also Figure S1.**

The top GWAS hit in the *MHA1* region residues in *AT1G62045*, which encodes a small ∼7 kDa protein of 67 amino acids. To examine whether AT1G62045 regulates *ACD6*-Est-1 activity, we overexpressed in the Ty-0 background the *MHA1* alleles of Ty-0 and Est-1 **(Figure 1C)**, which differ in two adjacent codons, with one of these causing a non-conservative change from asparagine to serine due to the top GWAS SNP **(Figure 1D)**. Overexpression of *MHA1*-Est-1 but not *MHA1*-Ty-0 caused strong leaf necrosis and reduced size in 13 out of 15 T_1_ individuals, supporting that *AT1G62045* is *MHA1* **(Figure 1C)**. *MHA1* homologs are found in many plant species **(Figure S1F)**, and alignment of the predicted protein sequences told us that other Brassicaceae typically encode the asparagine corresponding to the Est-1 variant, which is representative of the major *A. thaliana* allele **(Figure 1D)**.

The Ty-0 allele of *MHA1* carried a nonsynonymous substitution that affected a conserved asparagine **(Figure 1D)**, suggesting that this allele might have reduced or no activity, in agreement with the absence of effects in the overexpresssion experiment. Knockout of *MHA1*, however, led to cell death and increased *PR1* expression in Ty-0, indicating that *MHA1*-Ty-0 was functional **(Figure 1E)**. This was surprising, because the *MHA1*-Est-1 allele seemed also to be functional, since overexpression of *MHA1*-Est-1 in Ty-0 also caused cell death and reduced growth **(Figure 1C)**. Conversely, introduction of a genomic copy of *MHA1*-Ty-0 into Est- 1 *mha1* mutants suppressed cell death and increased biomass, in agreement with this allele interfering with the action of *ACD6*-Est-1 **(Figure S1G)**. Together, these data pointed to *MHA1*-Ty-0 as a gain-of-function allele that acts as a negative regulator of *ACD6*-Est-1 activity in cell death promotion. The standard allele, *MHA1*-Est-1, apparently masks the action of *MHA1*-Ty-0, but it is not required for *ACD6*-Est-1 activity.

### Genetic characterization of the *MHA1* paralog *MHA1L*

In the *A. thaliana* reference genome, *MHA1* and the adjacent gene *AT1G62050* are paralogous to the 3’ and 5’ portions of another gene on chromosome 1, *AT1G11740*. This gene model appears to be mis-annotated, as public RNA-seq data and RT-PCR analyses indicated that two genes are transcribed from AT1G11740 **(Figures S1H-S1K)**. We thus designated the *MHA1* paralog *MHA1-LIKE* (*MHA1L*). The gene duplication that gave rise to *MHA1* and *MHA1L* appears to have occurred at the base of the Brassicaceae as part of a whole-genome duplication^24^, and independent duplications are found in other lineages **(Figure S1F)**. A *mha1 mha1L* double mutant in Est-1 resembles the wild type plants in ADC6-mediated cell death, suggesting that neither *MHA1* nor *MHA1L* is required for *ACD6*-Est-1 activity **(Figure S1L)**.

As overexpression of *MHA1-*Est-1 constitutively activates immune responses, we further characterize the roles of MHA1 and MHAL1 in plant immunity. An expression atlas^25^ revealed that *MHA1* expression is elevated when plants are infected with the bacterium *Pseudomonas syringae* pv. *tomato* (*Pst*) DC3000 expressing *avrRPM1* or with the fungus *Botrytis cinerea. MHA1L* expression was also induced by *Pst* DC3000 infection (**Figures S2A**). To gain further insights into MHA1 family function in immunity, we generated *mha1*, *mha1l*, and *mha1 mha1l* double mutants in the Col-0 reference background, which has the standard *ACD6* allele (**Figures S2B**). Reactive oxygen species (ROS) burst and MAPK activation after flg22 treatment were weaker in *mha1l* and *acd6-*2 null mutants than in wild type **(Figures 2A, 2B, and S2C)**, suggesting that *MHA1L*, like *ACD6*, is involved in PTI. In agreement, *mha1l, mha1 mha1l*, and *acd6-*2 mutants were similarly hyper-susceptible to *Pst* DC3000 and *Pst* DC3000 *hrcC^-^* strain compared to Col-0 plants **(Figures 2C and 2D)**. However, neither *mha1l* nor *mha1 mha1l* had obvious defects in ETI **(Figures S2D and S2E).** There was a small change in ROS burst and MAPK activation, but not in *Pst* DC3000 growth in *mha1* single mutants **(Figures 2A-2D and S2C-S2E)**, suggesting that *MHA1* does not play a major role in defense in the presence of the *ACD6* reference allele in Col-0.

**Figure 2.**
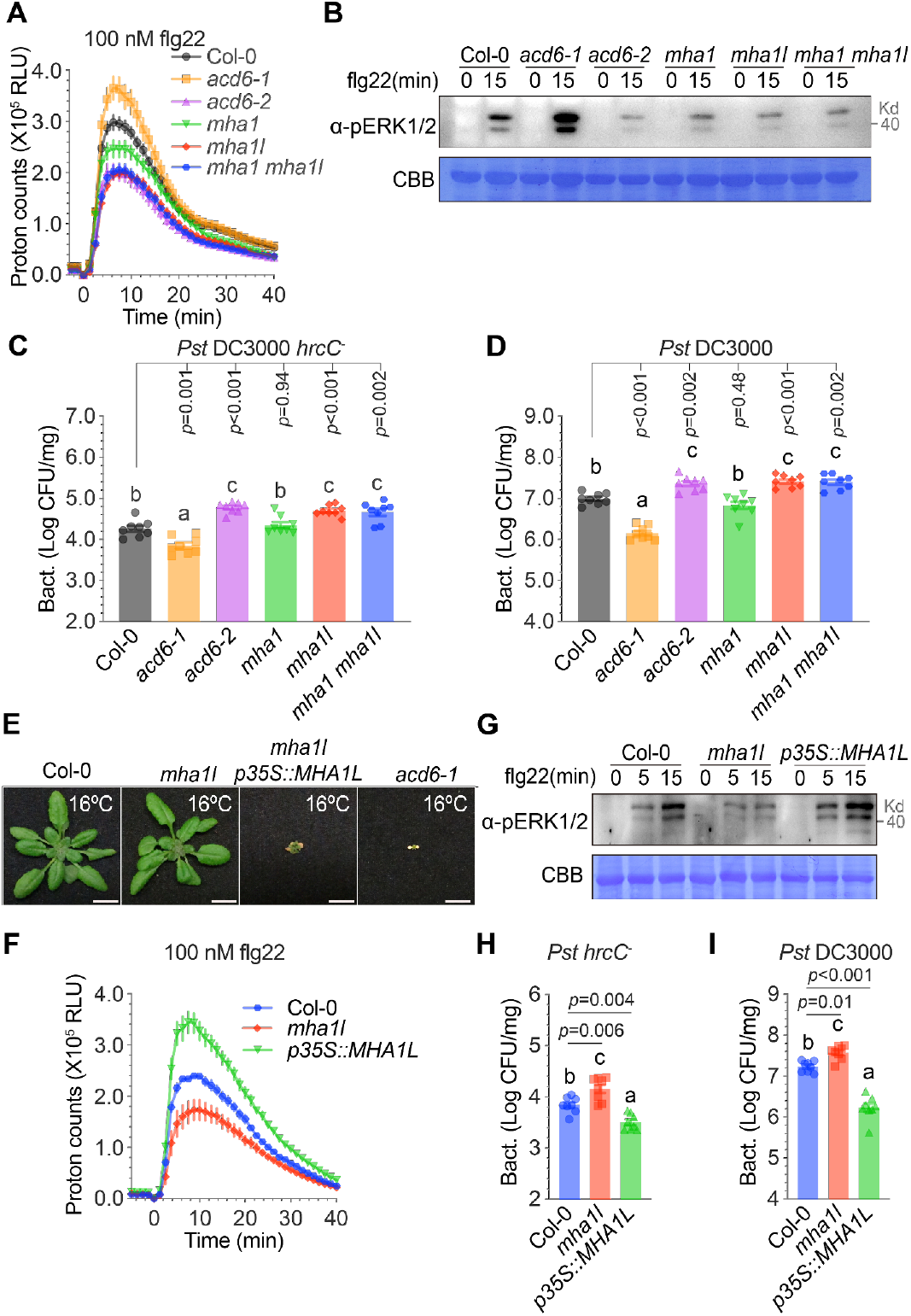
MHA1 and MHA1L are required for PTI and basal resistance. The *acd6*-1 gain-of-function and *acd6*-2 loss-of-function alleles were used as positive and negative controls. **(A, F)** Time course of flg22-induced ROS. Data are represented as mean ± s.e.m.; n = 12. See also Figure S2C. **(B, G)** flg22 induced MAPK activation, as detected with anti-pERK1/2. **(C, D, H, I)** Bacterial growth of *Pst* DC3000 *hrcC^-^*(C, H), and *Pst* DC3000 (D, I) 3 days after flood inoculation. Data are represented as mean ± s.e.m.; n = 8. *p*-values from Tukey’s HSD test. Letters indicate significantly different groups, as determined by one-way ANOVA followed by Tukey’s test at α = 0.05. **(E)** Photographs of 4-week-old Col-0, *mha1l*, *mha1l p35S::MHA1L* and *acd6*-1 plants at 16°C. Plants in the other panels were grown at 23°C. **See also Figure S2.**

We also compared the effects of increased *MHA1* and *MHA1L* activity in the reference accession Col-0. Neither *MHA1*-Est-1 nor *MHA1*-Ty-0 overexpression had obvious effects in Col-0, although *MHA1*-Est-1 had slightly higher *PR1* expression **(Figure S2F)**. In contrast, overexpression of *MHA1L*, which does not show allelic variation in amino acid sequence among Col-0, Est-1, and Ty-0, caused strong cell death and dwarfing at 16°C in 24 out of 30 T_1_ plants **(Figure S2G)**. The effects at 23°C were milder **(Figure S2G)**, which can be at least partially attributed to reduced accumulation of the protein at the higher temperature, as demonstrated with plants overexpressing an *GFP*-*MHA1L* fusion **(Figure S2H)**. The temperature sensitivity of the *MHA1L* overexpressors suggested that *MHA1L* can enhance *ACD6* activity, since it resembled the behavior of *acd6*-1 gain-of-function mutants, which also show enhanced cell death at 16°C compared to the standard temperature of 23°C **(Figures 2E and S2E)**^2^. Similarly, *MHA1L* overexpressors in *mha1l* showed enhanced ROS production, MAPK activation and resistance to *Pst* DC3000 and *Pst* DC3000 *hrcC^-^* bacteria **(Figures 2E-2I)**, in support of a role of *MHA1L* in PTI. Genetic analysis confirmed that *MHA1L* overexpression defects were dependent on SA, as the cell death phenotype was either partially or completely suppressed by mutations in key SA signaling components such as *NPR1* and helper NLRs of the ADR1 and NRG1 families **(Figures S3A-S3C)**.

### Identification of *ACD6* as a potent suppressor of *MHA1L* overexpression

To illuminate the role of MHA1L in immunity, we generated a mutagenized population of *MHA1L* overexpression in an unbiased manner, we mutagenized *MHA1L* overexpressors in the Col-0 background with ethyl methanesulfonate and screened for suppressors **(Figure 3A)**. In total, revealed that 37 out of the 72 suppressor lines carry mutations in ACD6 **(Figure 3B)**. Four mutations in *ACD6* were reported to be revertants of the *acd6*-1 gain-of-function allele^26^, consistent with them being loss-of-function alleles. Thirteen mutations affected residues conserved between ACD6 and its wheat homolog Lr14a, including L300F, which corresponds to L362F in Lr14a, which is a loss-of-function mutation in wheat^15^ **(Figure 3B,)**. In addition, co-segregation analysis showed that the *ACD6* locus co-segregated with suppression of *MHA1L*-associated cell death for one of *somha1l* lines **(Figure S3D)**. Final validation for *ACD6* being essential for the effects of *MHA1L* overexpression was obtained with the known *acd6*-2 loss-of-function allele, which suppressed the *MHA1L* overexpression phenotype as well (25 T_1_ individuals) **(Figure 3C)**. In contrast, the *acd6*-1 gain-of-function mutation greatly enhanced the phenotype of *MHA1L* overexpressors (15 T_1_ individuals), and *acd6*-1 *p35S::MHA1L* plants grown at 23°C resembled *MHA1L* overexpressors without the *acd6*-1 allele grown at 16°C **(Figure 3E)**.

**Figure 3.**
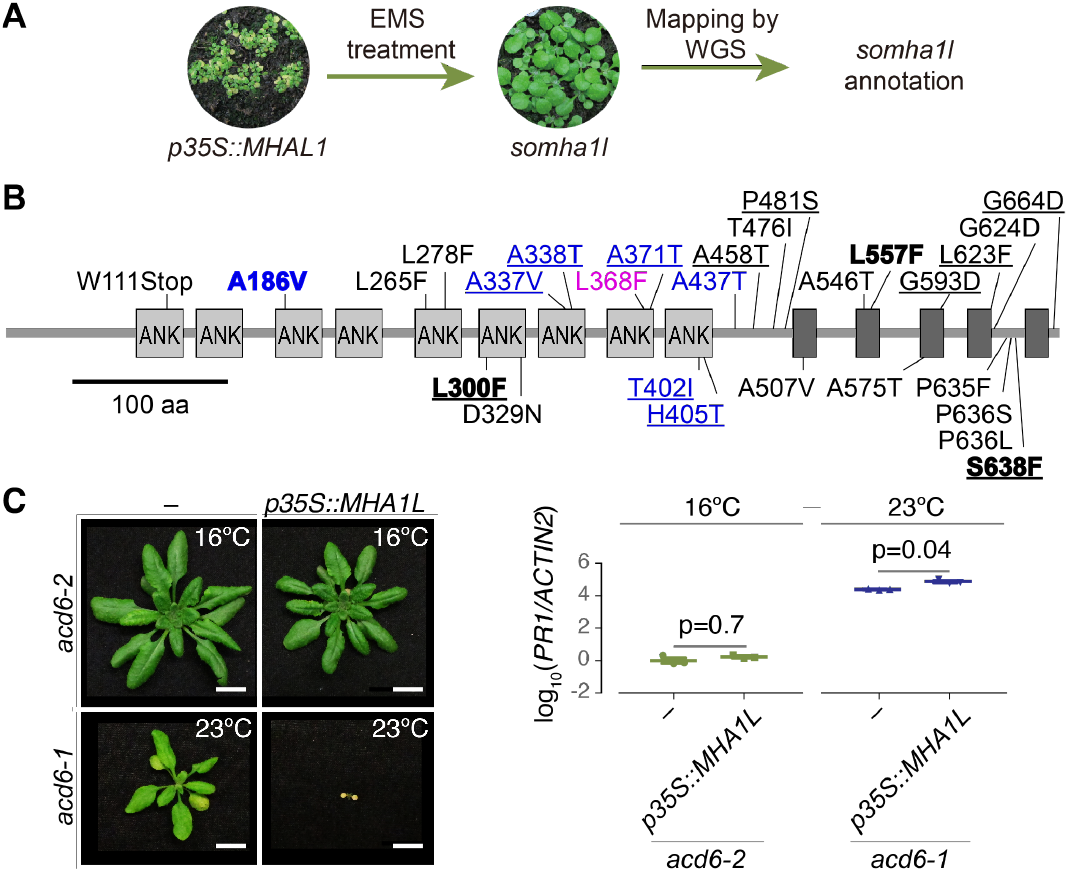
Identification of Suppressor of MHA1L. **(A)** Isolation of *suppressor of mha1l* (*somha1l*) mutants. **(B)** Map of 28 EMS-induced *somha1l* mutations in *ACD6* identified as suppressors of *MHA1L*-induced cell death. Numbers indicate codons. Four mutations in bold are identical to ones that have been isolated before as intragenic suppressors of the *acd6*-1 gain-of function allele^26^. Blue residues are identical in two out of three mammalian TRP proteins (TRPA1, NOMPC, TRPV6) with related ankyrin (ANK) repeats, magenta residue in all three. Residues identical in Lr14a^15^ underlined. redicted transmembrane helices shown as black boxes. **(C)** *acd6*-2 and *acd6*-1 plants overexpressing *MHA1L*. Left, plant phenotypes. Right, accumulation of *PR1* mRNA, as measured by qRT-PCR from three biological replicates each. *p*-values from Tukey’s HSD test. Scale bars 1 cm. **See also Figure S3, and Table S1.**

Together, the experiments with *MHA1L* overexpressors and *acd6* loss- and gain-of-function alleles in the Col-0 background indicated that *MHA1L* function depends on *ACD6* activity. However, not only was the *ACD6*-Est-1 phenotype not enhanced by *MHA1L* overexpression, but the severe *MHA1L* overexpression phenotype at 16°C was suppressed in Est-1 and in a NIL with the *ACD6*-Est-1 allele **(Figures S3E and S3F)**, indicating genetic interaction of *MHA1L* with the standard *ACD6*-Col-0, but not the *ACD6-*Est-1 allele **(Figure S2G)**. Note that the phenotype of *ACD6*-Est-1, in contrast to that of *acd6*-1, is not enhanced by lower temperature^1, 2^, which distinguishes *ACD6*-Est-1 from many other autoimmunity mutants^27, 28^. The genetic interactions and temperature effects between *MHA1/MHA1L* and *ACD6* alleles are summarized in **Figures S3G.**

### Characterization of interaction between ACD6 and MHA1L

The genetic reliance of *MHA1L* on *ACD6*-Col-0 in the Col-0 accession prompted us to test whether their encoding proteins interact. MHA1L interacts with both full length and the cytoplasmic ankyrin repeat fragment of ACD6 in split-luciferase complementation assays in *Nicotiana benthamiana* and co-immunoprecipitation (co-IP) in *A. thaliana* protoplasts **(Figures 4A-4C)**. Direct interaction between MHA1L and ACD6-Col-0 was further observed in the *in vitro* GST or MBP pull-down assays **(Figures 4D and 4E)**. In co-IP and GST pull-down assays **(Figures 4C and 4D)**, MHA1L tended to produce a stronger signal than MHA1, but more quantitative methods would be required to determine whether these apparent differences are significant.

**Figure 4.**
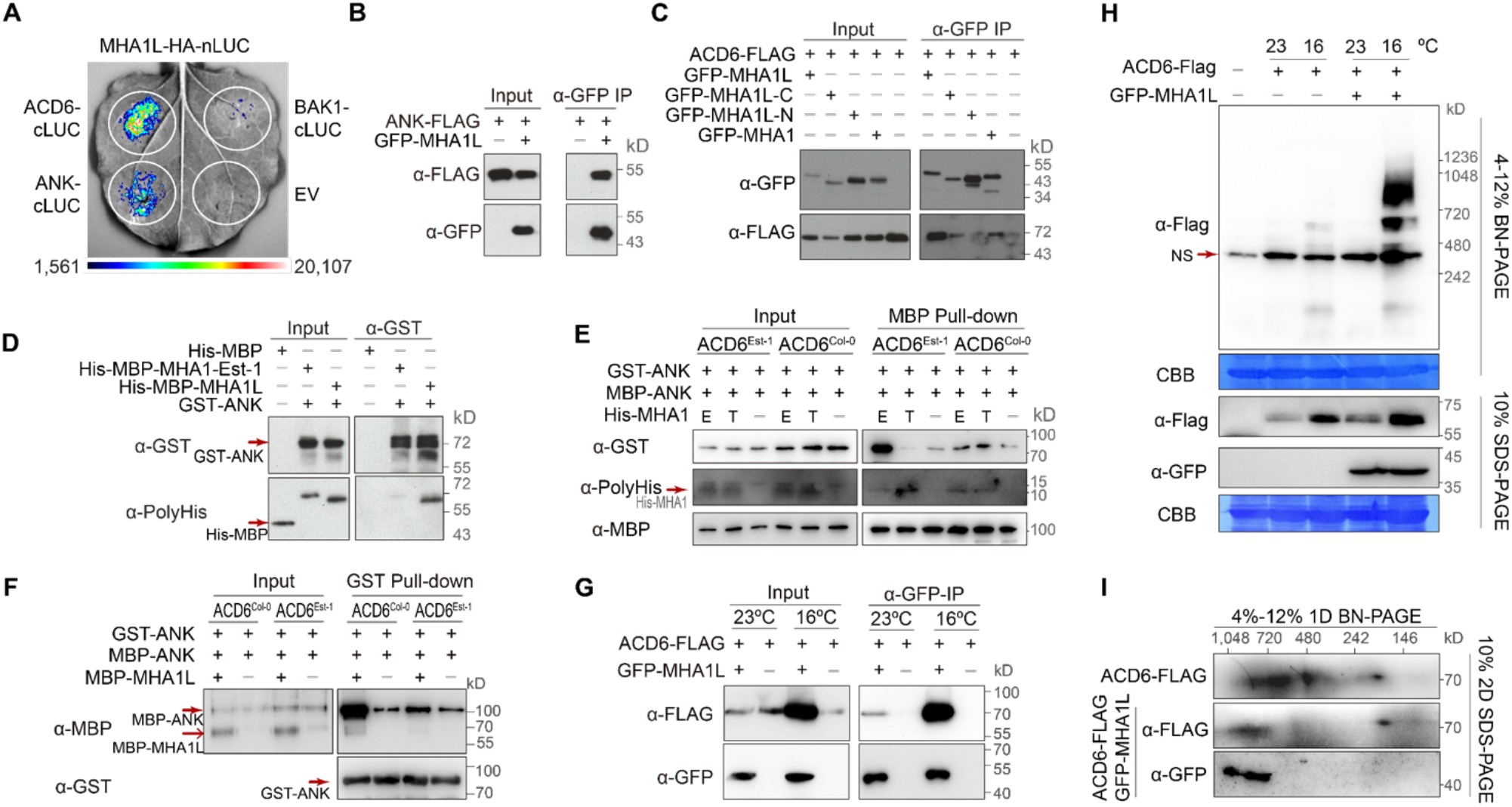
Low temperature enhances ACD6 accumulation and complex formation in MHA1L overexpressors. **(A)** Split-luciferase complementation assays of interaction of MHA1L with ACD6 from Col-0 and a cytoplasmic ACD6-ANK fragment in *Nicotiana benthamiana*. BAK1 served as a negative control. **(B)** Co-IP assays with extract from *A. thaliana* Col-0 protoplasts to probe interactions of ACD6 with MHA1L full-length protein and a C-terminal MHA1L fragment (residues 20-72). **(C)** Co-IP assays with extracts from *A. thaliana* protoplasts to probe *in vivo* interaction of a cytoplasmic ACD6-ANK fragment with MHA1L. **(D)** GST pull-down assay to probe *in vitro* interaction of the ACD6-ANK fragments with MHA1 and MHA1L. **(E)** MBP pull-down assay to probe *in vitro* ACD6-ANK self-association with or without MHA1s. E, Est-1; T, Ty-0. **(F)** GST pull-down assay to probe ACD6-ANK self-association with or without MHA1L. **(G)** Interaction of MHA1L with ACD6 *in vivo*, as shown by co-IP with material from transgenic *A. thaliana* plants. ACD6-FLAG was expressed from its native promoter and MHA1L expression was driven by the CaMV35S promoter. Seedlings were grown in 23°C short days for 12 days, or in 23°C for 10 days, then moved to 16°C for 2 days. **(H)** BN-PAGE and SDS-PAGE showing reduced ACD6 accumulation and complex formation in MHA1L overexpressors grown in 23°C compared to 16°C. Seedlings were grown in 23°C long days for 12 days, or for 10 days, then transferred to 16°C for 2 days. NS indicates nonspecific bands. **(I)** Evidence for MHA1L and ACD6 residing in the same complex, as shown with 2D SDS-PAGE. **See also Figure S4.**

Finally, we examined the subcellular localisation of MHA1 and MHA1L fused with GFP tag in Col-0. In both leaf and root cells, GFP signal was detected both in the cytoplasm and at the plasma membrane **(Figures S4A-S4D)**. Application of endocytosis inhibitor brefeldin A (BFA) enables GFP-MHA1 to co-localize endocytic tracer FM4-64 in the BFA bodies, suggesting that GFP-MHA1 recycles between the plasma membrane and endosomal compartments, similar to many other plant plasma-membrane proteins^29^ **(Figure S4B)**. Subcellular fractionation indicated that both GFP-MHA1 and GFP-MHA1L proteins were enriched in microsomes **(Figures S4E and S4F)**, similar to ACD6-1 protein^11^ **(Figure S4G)**. The similar subcellular localization of MHA1, MHA1L and ACD6 was in agreement with direct interaction of the proteins.

To better understand the genetic interactions, in which *MHA1L* overexpression did not affect *ACD6*-Est-1 activity, while *MHA1*-Ty-0 suppressed *ACD6*-Est-1 effects, we performed pull-down experiments to test how MHA1 variants affect self-association of ANK domains of ACD6. We found that MHA1-Est-1 greatly enhanced ANK-ACD6-Est-1 self-association. In contrast, MHA1-Ty-0 slightly inhibited ANK-ACD6-Est-1 self-association **(Figure 4E)**. Moreover, MHA1L enhanced the self-association of ANK-ACD6-Col-0 but not of ANK-ACD6-Est-1 **(Figure 4F)**, which is consistent with MHA1L overexpression not modifying the extent of ACD6-dependent leaf necrosis in Est-1.

We developed the following hypothesis for the MHA1/MHA1L interaction with ACD6: First, activity of the standard form of ACD6 (as found in Col-0) is enhanced by MHA1L; such an enhanced activity could provide additional feedforward regulation of *ACD6* in response to pathogen challenge, as both enhanced *ACD6* activity and infection of plants with a bacterial pathogen increased *MHA1L* mRNA accumulation **(Figure S3A)**. Second, the amino acid substitutions in the transmembrane portion that are causal for increased activity of ACD6-Est-1^1^ alter the conformation of ACD6-Est-1, such that it no longer requires binding of MHA1L to its ankyrin repeats for increased activity. ACD6-Est-1 can, however, be bound by MHA1-Ty-0, which in turn interferes with ACD6-Est-1 activity.

Since MHA1L overexpression had more drastic phenotypic effects at 16°C than at 23°C **(Figures 3B and S3B-S3F)**, we performed co-IP experiments at both temperatures. We found that higher temperature not only reduced ACD6 accumulation, but also ACD6 interaction with MHA1L **(Figure 4G)**. We further examined ACD6 accumulation in *A. thaliana* plants using Blue Native polyacrylamide gel electrophoresis (BN-PAGE), which has been previously used to show that ACD6 exists in complexes of around 700-800 kDa^11, 16^. Overexpression of MHA1L increased the total levels of ACD6 and that of large ACD6 complexes, which was also enhanced by lower temperature **(Figure 4H)**, similar to what has been shown for the mutation in *acd6*-1, which, like MHA1L overexpression, enhances ACD6 activity^10, 11^. Finally, analysis of complexes with 2D SDS-PAGE confirmed that MHA1L interacts with ACD6 and promotes ACD6 complex formation, with MHA1L and ACD6 being present in high-molecular-mass complexes of similar sizes **(Figure 4I).**

### MHA1L enhances ACD6-stimulated ion channel activity

Genetic and biochemical experiments had indicated that MHA1L strongly modulates ACD6 activity. To glean more clues about ACD6’s function, we performed HHpred profile Hidden Markov Model searches^30^. We found multiple excellent hits to transient receptor potential (TRP) channels from animals and fungi, which regulate ion flux in response to stimuli ranging from heat to natural products and proinflammatory agents^31, 32^ **(Figure S5A)**. Alignments revealed extensive similarity between the ankyrin repeats of ACD6 and TRP proteins **(Figure S5B)**, but not between their transmembrane domains^33^. Out of 12 loss-of-function mutations mapping to the ACD6 ankyrin repeats, eight of the affected residues were identical in alignments to at least two out of the three top HHPred hits, fly NOMPC, human TRPA1, and rat TRPV6 **(Figure 3B)**. The *A. thaliana* genome encodes over 100 ankyrin repeat proteins, with many of them having transmembrane domains^34^. We could reliably identify five transmembrane domains in ACD6^35^. In contrast, TRP channels have six transmembrane helices, with helices 5 and 6 forming the ion pore in a functional TRP multimeric ion channel^31, 32^. Homology modeling did not suggest that the structure of the ACD6 transmembrane domains is consistent with those of TRP channels **(Figures S5C and S5D)**.

Although the similarities between TRP and ACD6 were restricted to the ankyrin repeats, ACD6 has multiple transmembrane domains that might form a pore, and we therefore investigated potential channel activity of ACD6 using African clawed frog (*Xenopus laevis*) oocytes. Since calcium has been implicated in different steps of immune signaling^36–38^, we were particularly interested in potential ACD6 activity as a calcium channel or regulator of calcium channels. Unfortunately, direct observation of calcium influx is hampered in *Xenopus* oocytes by the presence of endogenous calcium-activated chloride channels, which are stimulated by the accumulation of intracellular calcium and typically mask small ionic currents from calcium influx. However, these endogenous channels can be used as quantitative readout for channel-mediated calcium influx by foreign functional calcium channels^20, 39^.

When we raised the external calcium concentration in our experiments with oocytes expressing standard ACD6 from Col-0 or the gain-of-function ACD6-1 variant, we observed ionic currents that resembled calcium-activated chloride currents in their current-voltage relationship; these currents were lost upon removal of external chloride **(Figures 5A and 5B)**. The calcium-activated chloride currents were much lower upon expression of the ACD6^L557F^ variant, which has a substitution in the transmembrane domain **(Figure 5C)**. Consistent with the role of MHA1L as an ACD6 activator in plants, MHA1L could further enhance the currents in ACD6-expressing oocytes **(Figure 5D)**. This was not the case when MHA1L was co-expressed with ACD6^L300F^, which has a substitution in the ankyrin domain **(Figures 5C and 5D)**.

**Figure 5.**
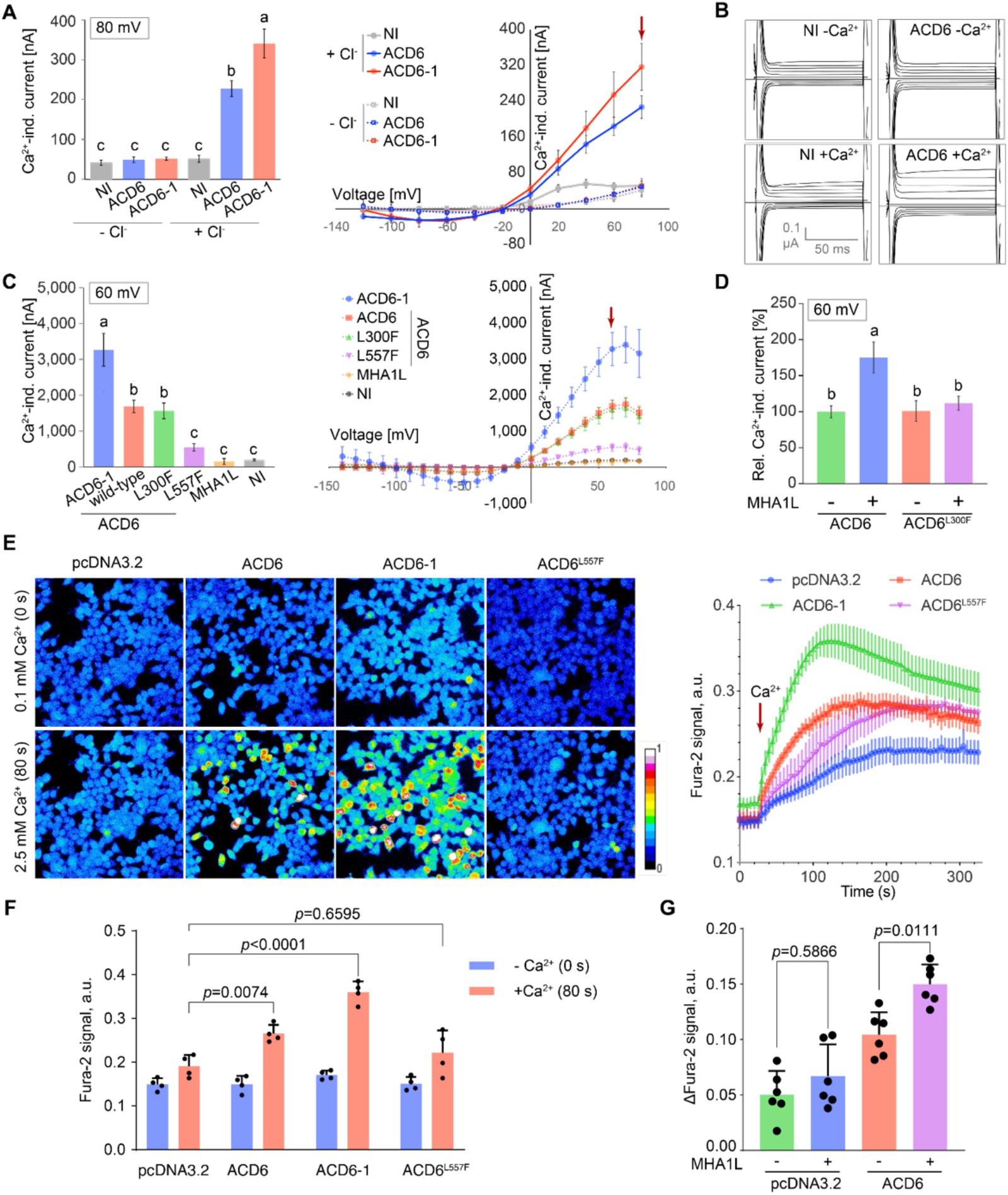
Stimulation of ion channel activity by ACD6and enhanced stimulation in the presence of MHA1L in *Xenopus* oocytes and HEK293 cells. **(A)** Currents induced by 10 mM Ca^2+^ in *Xenopus* oocytes expressing ACD6 variants in the presence or absence of Cl^-^ at 80 mV. In the current-voltage plot on the right, the data points for ACD6-1 without Cl^-^ are obscured by the ACD6 data points without Cl^-^. NI, not injected (negative control). **(B)** Representative original current traces from ACD6 recordings in (A). **(C)** Left, ionic currents induced by 10 mM Ca^2+^ in *Xenopus* oocytes at +60 mV. L300F is a substitution in the ankyrin repeat domain, L557F is a substitution in the transmembrane region of ACD6. Shown are means (n≥4) ± SE, letters indicate significantly different groups (p<0.01; post hoc Tukey’s HSD). Right, ionic currents induced by 10 mM Ca^2+^ in *Xenopus* oocytes represented as a current-voltage plot of a representative oocyte batch. Shown are means (n≥8) ± SE. **(D)** Relative currents at +60 mV of three different oocyte batches, normalized to the mean of wild-type ACD6 of each batch. Means (n≥ 8) ±s.e.m., letters indicate significantly different groups (p<0.05; post hoc Tukey’s HSD). **(E)** Left, Fura-2 signal with different extracellular Ca^2+^ concentrations in HEK293 cells. pcDNA3.2 indicates cells transformed with an empty vector. Scale shown on left, 0 indicates minimum signal, 1 maximum signal. Right, time course of Fura-2 signal. Time point at which extracellular Ca^2+^ was increased from 0.1 to 2.5 mM is indicated by an arrow. Data are the means ± s.e.m (n=4; a.u., arbitrary units). **(F)** Increase in Fura-2 signal 80 s after extracellular Ca2+ concentration was raised from 0.1 mM to 2.5 mM. Same experiment as in **E**. Data are means ± s.e.m (n=4) analyzed by two-way ANOVA, *p*-values from Tukey’s honest significant difference (HSD). **(G)** Increase in Fura-2 signal in response to co-expression of MHA1L. Data are means ± s.e.m. (n=6) analyzed by one-way ANOVA, *p*-values from Tukey’s HSD. **See also Figure S5.**

To verify the observation in the oocyte system, we transiently expressed standard ACD6 from Col-0, the gain-of-function ACD6-1 variant or the ACD6^L557F^ variant in human embryonic kidney 293 (HEK293) cells. Codon optimization was required for efficient expression of ACD6 (“opti2-ACD6”) in HEK293 cells **(Figures S5E and S5F)**. opti2-ACD6-mGFP5-6xHis and MHA1-L-mGFP5-6xHis were both observed at the plasma membrane **(Figure S5F)**, with stronger GFP signal enriched at the plasma membrane of dying cells **(Figure S5G)**. Calcium influx induced by increasing extracellular calcium, from 0.1 mM to 2.5 mM, was monitored using the ratiometric fluorescent dye Fura-2, which binds to intracellular calcium^23, 40–42^. Compared to cells transformed with an empty vector, calcium influx was increased in cells expressing opti2-ACD6-myc-6xHis from Col-0 or the gain-of-function variant opti2-ACD6-1-myc-6xHis, but not in cells expressing the opti2-ACD6^L557F^-myc-6xHis loss-of-function variant **(Figures 5E and 5F)**. Co-expression of MHA1L-myc-6xHis enhanced the calcium influx in response to elevation of extracellular calcium, recapitulating the observations from oocytes **(Figure 5G)**. Together, these results show that ACD6 can stimulate ion channel activity in two different heterologous systems, *Xenopus* oocytes and human HEK293 cells, and that this activity can be further enhanced by MHA1L.

### MHA1L and ACD6 are required for calcium signaling in PTI responses

To obtain *in planta* evidence for ACD6 being involved in calcium influx, we introduced the luminescent aequorin calcium biosensor, which has been used to assay responses to pathogen- and damage-associated molecular patterns (PAMPs/DAMPs)^43–45^, into *acd6*-ko, *acd6*-1 and *MHA1L* overexpressor backgrounds, using the calcium channel mutant *death, no defense1* (*dnd1*-1) as a positive control^20^. Compared to wild-type plants, flg22-induced calcium influx was greatly reduced in *acd6*-ko loss-of-function mutants **(Figure 6A)**. We confirmed this finding with the R-GECO1 calcium reporter, which has also been used to monitor flg22-induced calcium fluxes^46^ **(Figure S6A)**. Similarly, loss-of-function of MHA1L impaired calcium influx triggered by flg22 treatment **(Figure 6A)**, which is consistent with impaired flg22-triggered ROS burst and MAPK activation in *mha1l* mutants **(Figures 2A and 2B)**. These were specific defects, since the PAMP chitin and the stressor NaCl did not affect calcium influxes in either *acd6*-ko or *mha1l* mutants **(Figures 6B and 6C)**. Similar specificity was observed when *ACD6*-Est-1 was knocked out in the Est-1 background **(Figures S6C-S6F)**. As expected, flg22-induced calcium influx was greatly increased by *MHA1L* overexpression **(Figure 6A)**. Together with the observation that MHA1L and ACD6 can stimulate calcium channel activity in *Xenopus* oocytes and human HEK293 cells, we propose that ACD6 contributes to the regulation of calcium influx, which can be directly activated by MHA1L during plant immunity.

**Figure 6.**
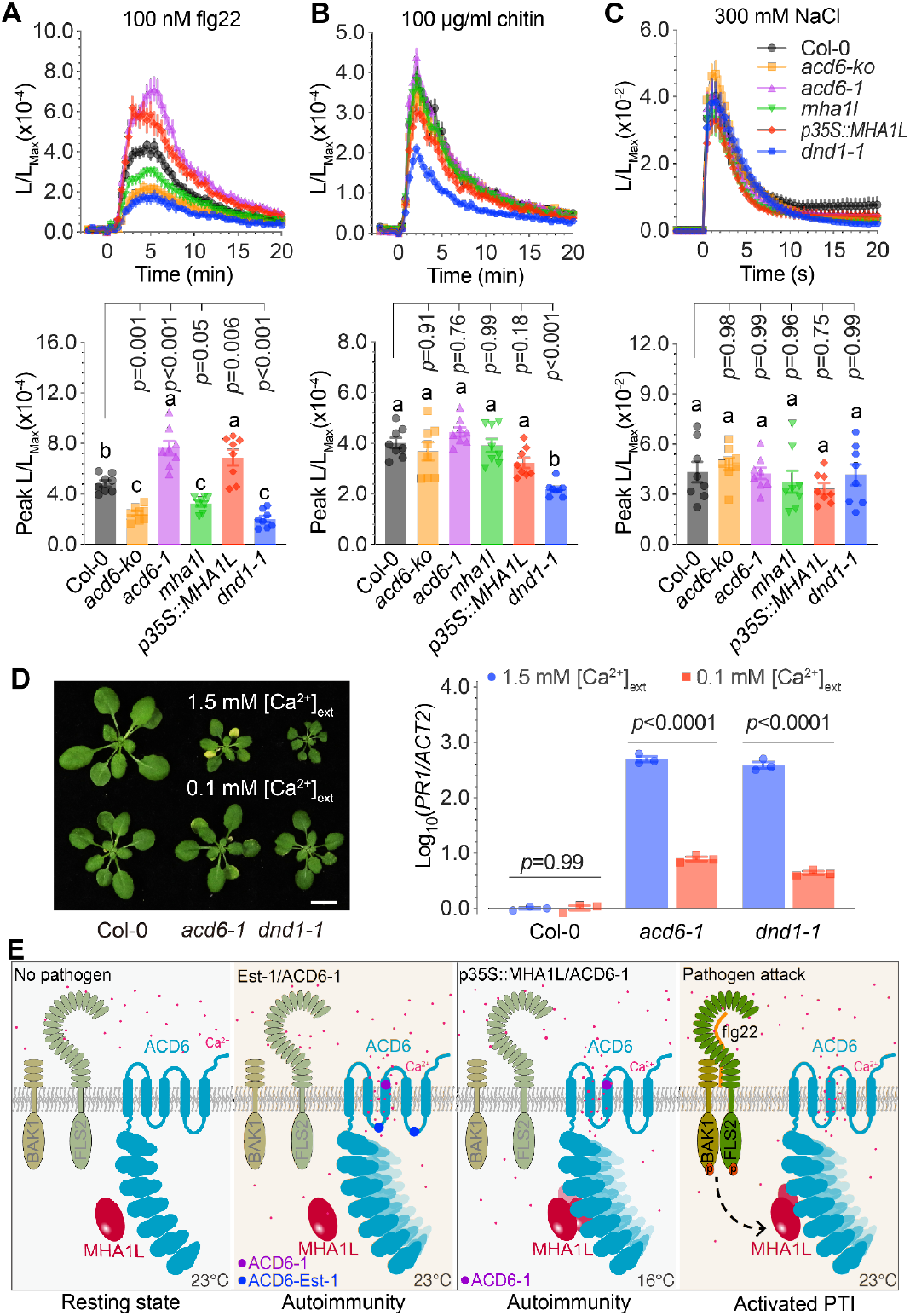
Requirements of *ACD6* and *MHA1L* for calcium signaling in PTI. **(A-C)** Top, time courses of aequorin-luminescence in 5-day-old seedlings after addition of 100 nM flg22 (**A**), 100 μg/ml chitin (**B**), or 300 mM NaCl (**C**). Bottom, peak values. Data are means ± s.e.m. (n=8), analyzed by one-way ANOVA, *p*-values from Tukey’s HSD. Letters indicate significantly different groups at α = 0.05. **(D)** Left, rosettes of 3-week-old plants grown with different concentrations of external Ca^2+^. Right, accumulation of *PR1* mRNA (n=3). The experiment was repeated twice, with similar results. Scale bar 1 cm. *p*-values from Student’s t-test. **(E)** Model for MHA1L-ACD6 regulation in immunity. Under normal conditions, MHA1l-ACD6-associated complexes are maintained at a low level. ACD6-Est-1 and *acd6-1* gain-of-function mutant show high ion channel activity independent of MHA1L, which results in autoimmunity. The formation of ACD6-associated complexes is constitutively promoted by MHA1L overexpression or *acd6-1* gain-of-function mutant to ion channel activity in low temperature, and results in autoimmunity. When plants are attacked by pathogens, PAMPs-PRRs interaction triggers formation of MHA1L-ACD6-associated complexes to promote immune responses. **See also Figure S6**.

Since ACD6 could stimulate calcium influx in both animal and plant cells, we further tested whether calcium is required for *ACD6-*triggered immunity. To this end, we cultured the gain-of-function mutant *acd6*-1 in a hydroponic system that contained either 1.5 or 0.1 mM external calcium. Similar to the calcium channel mutant *dnd1-*1^20^, we found that depletion of calcium attenuated signs of autoimmunity and expression of *PR1* in *acd6*-1 **(Figure 6D)**.

## DISCUSSION

Starting with *A. thaliana* accessions carrying the naturally highly active *ACD6*-Est-1 allele, whose phenotypic effects in different genetic backgrounds depend on unlinked modifiers, we identified a new gene family encoding small protein regulators of plant immunity. Follow-up genetic and biochemical experiments revealed not only complex interactions between different natural variants of *MHA1*, *MHA1L* and *ACD6*, but also that MHA1 and MHA1L are direct regulators of ACD6 activity by binding to the N-terminal ankyrin repeats of the ACD6 transmembrane protein.

The ankyrin repeats of ACD6 are similar in sequence to those of mammalian TRP proteins, and similar to certain TRP proteins, ACD6 functions can stimulate ion flux across membranes **(Figure S5)**. There is extensive knowledge on the structure and function of TRP proteins, providing potential models for understanding the activity of ACD6^31–33^. However, the transmembrane domains of ACD6 and TRP channels differ, and it is therefore unlikely that ACD6 and homologs function in exactly the same manner as TRP proteins do. Similar to a subset of TRP proteins, ACD6 can apparently be regulated through its ankyrin repeats by small ligands, the MHA1 and MHA1L proteins, with MHA1L enhancing the accumulation and activity of both the standard ACD6 protein and its experimentally induced derivative ACD6-1, but not of the natural ACD6-Est-1 variant. In contrast, MHA1-Ty-0, but not the standard MHA1 protein, can suppress activity of the ACD6-Est-1 variant **(Figures 6E and S3G)**.

TRP channels typically function as tetramers, with two membrane-spanning helices from each subunit contributing to the ion channel pore (see Ref. 33 for a recent compilation of TRP channel structures). ACD6 is also found in a large complex^11, 16^ **(Figures 4H and 4I)**, but the size of this complex is substantially greater than that of a simple tetramer, suggesting either a higher oligomeric state or association with additional proteins. Several TRP channels can form heteromultimers, with heteromultimers often having different activities than the homomultimers^31^. In *A. thaliana*, combinations of natural *ACD6* alleles, including those from the Mir-0 and Se-0 accessions, can lead to increased ACD6 activity in F_1_ hybrids relative to either parent^2, 3^, compatible with a scenario in which the variants assemble into heteromultimers in the F_1_ hybrids that have different properties than the respective homomultimers.

Also similar to *ACD6*, gain-of-function mutations in TRP channel genes are common, with many of these having been identified as variants underlying human genetic diseases^47^. *ACD6* itself was originally discovered based on the experimentally induced gain-of-function, hyperactive *acd6*-1 allele^4^. A highly active natural allele, *ACD6*-Est-1, was subsequently found to be common in natural populations of *A. thaliana*^1^. The Mir-0/Se-0 combination described above can also be considered as a gain-of-function situation. In all four alleles, the sequences responsible for increased activity are found in the predicted transmembrane portion of ACD6 **(Figure S6G)**. While some gain-of-function mutations in TRP channels affect either the N-terminal portions including the ankyrin repeats or the C-terminal portions, many alter amino acids in the transmembrane domain **(Table S1)**. This is significant, because, as in ACD6, the overall length of the transmembrane domains is typically much shorter than the length of the other domains, making for a smaller mutational target.

TRP channels can integrate inputs consisting of different stimuli or ligands, and the opposite effects of MHA1 and MHA1L on ACD6 activity are reminiscent of the effects of different ligands on TRPV1 activity^48^. The exact role of MHA1 in regulating ACD6 remains somewhat enigmatic, because the clearest effects are exerted by a natural dominant-negative allele that attenuates the activity of the highly active *ACD6*-Est-1 allele. It is conceivable that MHA1 is primarily a regulator of an ACD6 homolog; *A. thaliana* genomes encode about half a dozen proteins with close similarity to ACD6 in both the ankyrin repeats and transmembrane domains. Redundancy among these proteins may also explain why inactivation of *ACD6* has relatively minor consequences^10^ compared to the pronounced phenotypic effects resulting from *ACD6* hyperactivity or MHA1L overexpression (refs. ^1–4^ and this work). On the other hand, the effects of MHA1L overexpression seem to depend entirely on ACD6, pointing to an exclusive relationship between MHA1L and ACD6. It may appear surprising that MHA1L enhances accumulation of an ACD6 complex, rather than merely regulating its activity, but increased accumulation of ACD6 has also been observed in *acd6*-1 mutants^11^, consistent with positive feedback regulation of ACD6. One possibility is that MHA1L affects ACD6 activity at least in part by stabilizing ACD6, which has been shown to be regulated by protein degradation in the cytoplasm^11^, via other membrane-resident proteins with which ACD6 associates^11, 12, 16^.

Although the ankyrin repeats of TRP proteins are similar to those of ACD6 and related plant proteins, it is unlikely that the ACD6 transmembrane domains adopt a topology similar to those of TRP channels and it is therefore not obvious from the secondary structure whether ACD6 acts as an ion channel **(Figure S5C)**. Nevertheless, ACD6 can stimulate calcium influx both in two heterologous systems, *Xenopus* oocytes and human cell culture, and in plants, and several *ACD6* effects *in planta* appear to require calcium **(Figures 5 and 6)**. Calcium influx from extracellular stores is an early event in the host response to pathogens, with relatively well characterized downstream steps. One of the earliest cellular responses to PAMPs is the rapid increase of cytoplasmic calcium levels, indicating that calcium channels are intimately associated with PTI. In agreement, genetics has implicated cyclic nucleotide-gated channel (CNGCs) in this process, and at least one of these CNGCs is a functional calcium channel whose activity is regulated by PAMPs/DAMPs, such as bacterial lipopolysaccharide (LPS) and flg22 or plant elicitor peptide 3 (Pep3)^20, 36–38, 49, 50^. PAMPs also regulate activity of calcium channels from the OSCA family, which are important in stomata-dependent immunity^21^.

Similar to the experimentally induced *acd6*-1 and the natural *ACD6*-Est-1 alleles, mutations in genes for several plant calcium channels and transporters, including the ACA1 and ACA4 calcium-ATPases, the CAX1 and CAX3 vacuolar H^+^/calcium transporters, and the CNGC2/DND1 and CNGC4/DND2 calcium channels, can cause autoimmunity^20, 51–53^. We propose that ACD6 modifies PAMP-triggered calcium signals, supported by the finding that the calcium environment affects both autoimmunity in *acd6-*1 gain-of-function mutants and the proliferation of PTI-inducing *Pst* DC3000 *hrcC^-^* bacteria in *acd6-*2 loss-of-function mutants **(Figure 2C)**. Calcium also plays a role in ETI, as *cngc2*/*dnd1* mutants are impaired in the response to both avirulent and virulent bacterial pathogens^51^. Moreover, calcium channel blockers suppress cell death activated by the NLRs RPM1 and ZAR1^54, 55^, and ETI has recently been linked directly to calcium influx through the discovery that diverse NLRs and hNLRs are calcium permeable channels themselves^22, 23^. *ACD6* in turn has been genetically linked to the NLR SNC1^17^ and to helper NLRs of the ADR1 and NRG1 families **(Figure S3C)**. Our discovery of the MHA1/MHA1L family of peptides strengthens the case for ACD6 being a dynamic immune regulator associated with calcium influx **(Figure S6G)**, since MHA1L, which can activate strong, ACD6-dependent immune responses, appears to be transcriptionally induced upon pathogen infection **(Figure S2A)**. In addition, it has been reported that ACD6 and FLS2 form complexes in planta, with plasma membrane association of both FLS2 and BAK1 in response to SA signaling being enhanced by ACD6^11^. These observations are in agreement with our finding that ACD6 has an important role in PTI, regulating calcium influx in response to the PAMP flg22.

Several other ACD6-related transmembrane domain proteins with ankyrin repeats are involved in plant immunity. Cereals encode a series of ACD6-related proteins^56^. One of these has been implicated in resistance to the smut fungus *Ustilago maydis* in maize^14^, while another one, *Lr14a*, has been shown directly to confer rust resistance in wheat^15^. Many of the residues that are required for ACD6 function are conserved in Lr14a **(Figure 3B)**. In *A. thaliana*, BDA1, predicted to have four instead of five transmembrane segments as in ACD6, is required for activity of the receptor-like protein SUPPRESSOR OF NPR1-1, CONSTITUTIVE 2 (SNC2) in plant immunity^13^. As with the induced *acd6*-1 gain-of-function allele and the naturally highly active *ACD6* alleles, a mutation causal for *BDA1* gain-of-function activity maps to the transmembrane domain. Whether any of these ACD6 homologs depend at least in part on small peptide ligands remains to be determined.

In summary, our results demonstrate a complex set of relationships between different alleles and paralogs in the ACD6/MHA1/MHA1L system of small peptide-modulated calcium influx during pathogen responses, as well as positive and negative gain-of-function activities. Further complexity is added by the aggregate activity in this system being either temperature-sensitive or -insensitive, depending on the *MHA1L*- requirement of the *ACD6* allele **(Figure S3G)**. Our findings thus illustrate once more how naturally evolved special alleles, which are unlikely to be recoverable from conventional mutant screens, can provide new insights into fundamental aspects of biology.

## METHODS

### Plant material and growth conditions

*Arabidopsis thaliana* accessions and *Nicotiana benthamiana* were derived from stocks maintained in the lab. Seeds were germinated and cultivated in growth rooms at a constant temperature of 23°C or 16°C, air humidity at 65%, 16 h (long days) or 8 h (short days) 110 to 140 µmol m^-2^ s^-1^ light provided by Philips GreenPower TLED modules (Philips Lighting GmbH, Hamburg, Germany) with a mixture of 2:1 DR/W LB (deep red/white mixture with ca. 15% blue) and W HB (white with ca. 25% blue), respectively.

The hydroponic system used for assessing the effects of calcium has been described^67^ and was used with minor modifications. In brief, the plants were cultivated on inorganic solid media, and all nutrients were provided through the watering solution. The hydroponic medium contained 1 mM KH_2_PO_4_, 1 mM MgSO_4_, 0.25 mM K_2_SO_4_, 0.1 mM C_10_H_12_FeN_2_NaO_8_, micronutrients (50 µM KCl, 30 µM H_3_BO_3_, 5 µM MnSO_4_, 15 µM ZnSO_4_, 1 µM CuSO_4_, 1 µM NaMoO_4_), 1 mM NH_4_NO_3_, 1 mM NH_4_Cl, 1 mM KNO_3_ and different concentrations of CaCl_2_ (pH 5.8, adjusted with KOH).

### Genome-wide association study (GWAS)

Severity of cell death (leaf necrosis) was scored on an arbitrary scale from 1 to 5 using six biological replicates as described^17^, and GWAS with efficient mixed-model (EMMAX) methods was performed with the easyGWAS web interface^61^. The Bonferroni correction with a threshold of 0.05 for multiple testing corresponded to an uncorrected *p*-value of 7.6×10^-8^. The variance explained (adjusted R^2^) by the two loci was estimated using a Generalized Linear Model (GLM) using R^68^, with cell death as response variable and the two SNPs targeting the MHA loci as fixed effects with an interaction term.

### Linkage disequilibrium calculation

Genomic regions surrounding *MHA1* and *MHA2* were subset from a short read VCF of 1001 Genomes data^69^ using vcftools version 15.1 (ref. ^70^). Linkage disequilibrium R^2^ values were calculated with PLINK version 1.9, with a window of 15 kb and an R^2^ threshold of 0.

### Transgene-free genome-edited lines

An *A. thaliana* codon-optimized Cas9 (*athCas9*)^71^ was used, with the final *pUBQ10::athCas9:trbcs::gRNA1::gRNA2::mCherry* constructs assembled from six GreenGate modules^72^. Module A – *A. thaliana pUBQ10*; B – *A. thaliana* codon-optimized *Cas9*; C – *rbcs* terminator; D and E – sgRNAs expressed from *A. thaliana* U6 promoter^71^; The paired gRNA sequences for MHA1 are 5’- TAATAACATGTAGGCAACT-3’ and 5’-GGCATCTCCCCGATCCCTC-3’, and for MHA1L are 5’- GGAATTGGGGTGATGGCTG-3’ and 5’-GATGAAGAATCTTGTGCAG-3’, for ACD6 are 5’- GTGTCGCCCGTAGGTGACG-3’ and 5’-CGGTCCACGTTGTCGTCTG-3’ . F – *pAT2S3::mCherry:tMAS* cassette^73^ for selection of seeds with or without the transgene. Target regions were PCR amplified using oligonucleotide primers in **Table S2**.

### qRT-PCR

RNA was extracted from at least three biological replicates of pooled seedlings using the RNeasy kit (Thermo Fisher Scientific, Waltham, MA, USA), and treated with DNase I (Thermo Fisher Scientific). We used 2 µg of high-quality samples with A260/A230>2.0 and A260/A280>1.8 as determined with a ND-2000 spectrophotometer (Nanodrop Technologies, San Francisco CA, USA) as template for reverse transcription with the M-MLV Reverse Transcriptase kit (Thermo Fisher Scientific). Quantitative real-time PCR reactions were performed using Maxima SYBR Green Master Mix (Thermo Fisher Scientific) on a CFX384 instrument (Bio-Rad, Hercules, CA, USA). Transcript abundance was normalized to *ACTIN* 2 (*AT3G18780*). Primers are listed in **Table S2**.

### Transgenic lines

An *ACD6* fragment corresponding to Chr4: 8292084..8298360 in the TAIR10 reference genome was amplified from Ty-0 genomic DNA with PCR primers designed based on the *ACD6*-Est-1 sequence. To generate *pACD6::ACD6-3xFLAG* (*ACD6-FLAG*), genomic DNA was amplified from Est-1 and Col-0 plants using an oligonucleotide primer that contained a 3xFLAG coding sequence. Fragments were cloned into Gateway entry vector pCR8/GW/TOPO (Thermo Fisher Scientific) and moved into the binary vector pFK206, a modified pGREEN vector with *rbsc* terminator^74^ .

For *MHA1, MHA2* and *MHA1L* overexpression constructs, coding regions were amplified and cloned into pCR8/GW/TOPO, moved into binary vector pFK210, a modified pGREEN vector with a *35S* CaMV promoter and *rbsc* terminator^74^, and introduced into pFAST-G02 (ref. ^59^), which contains a seed coat fluorescence marker. To tag MHA1 and MHA1L proteins, mGFP coding sequences were added to *MHA1* and *MHA1L* coding sequences using Gibson assembly (NEB, Ipswich, MA, USA). We also generated 2.1 kb genomic constructs for both *MHA1*-Est-1 and *MHA1*-Ty-0 (*pMHA1::MHA1-Est-1* and *pMHA1::MHA1-Ty-0*) with pCR8/GW/TOPO and pFK206.

### Mutant screen for *p35S::MHA1L* suppressors

20,000 seeds of a *p35S::MHA1L* homozygous line were treated with 0.2% EMS overnight, followed by thorough rinses with water and sowing the seeds on soil. M_2_ seeds were collected in 640 pools. In the M_2_ generation, we screened for non-necrotic, normal-sized plants. Phenotypic suppression was confirmed in the M_3_ generation. Genomic DNA of M_3_ plants was extracted, tagmented^75, 76^, and sequenced on an HiSeq3000 instrument (Illumina, CA, USA) with 150 bp single-end reads. Raw reads were mapped to the TAIR10 reference genome using BWA-MEM^77^. bcftools^78^ and vcftools^70^ were used to call SNPs and generate the final VCF file, followed by snpEff^79^ annotation of SNP effects. Mutations shared by all lines were removed, and genes with at least three independent G>A or C>T mutations, typical for EMS mutations, predicted to cause non-synonymous changes or truncate the open reading frame were analyzed in detail.

### Trypan Blue staining

Freshly harvested leaf tissue was stained by completely immersing it in lacto-phenol/Trypan Blue staining solution (10 ml lactic acid, 10 ml glycerol, 10 ml phenol, 10 mg Trypan Blue and 10 ml water) and heating in a heat block at 80°C for 1 hour. Staining solution was aspirated and replaced by chloral hydrate solution (2.5 g/ml) to destain and clear the tissue. This was repeated once overnight to improve clearing. Samples were kept in 60% (v/v) aqueous glycerol for storage and further imaging.

### Measurement of ROS production

Leaf discs (6 mm in diameter) punched from rosette leaves of four-week-old soil-grown *A. thaliana* plants were incubated in 100 μl ddH_2_O in a 96-well white plate overnight. Water was replaced by 100 μl of a reaction solution containing 10 μM L-012 (Wako, Chuo-ku, Japan), and 10 μg/ml horseradish peroxidase (HRP, Solarbio Life Science, Beijing, China) supplemented with 100 nM flg22. Luminescence was measured with a microplate reader (BioTek Synergy H1, Agilent, CA, USA).

### MAPK activation

Two-week-old seedlings grown on ½ MS medium were treated with 100 nM flg22, then harvested and frozen in liquid nitrogen at indicated time. Total protein was extracted with extraction buffer containing 50 mM HEPEs-KOH (pH 7.5), 150 mM KCl, 1 mM EDTA, 0.2% Triton-X 100, 1 mM DTT, complete protease inhibitors and phosphatase inhibitors (Roche) after grinding in liquid nitrogen. Proteins were detected by immunoblotting using anti-phospho ERK1/2 antibodies (Cell Signaling Technology, MA, USA).

### Pathogen infection

Flood-inoculation and syringe-inoculation were used in this study. For flood-inoculation, three-week-old seedlings grown on ½ MS plate in short days were inoculated by *Pst* DC3000 or *Pst hrcC*^-^ as described^80^. Bacteria grown on King’s B (KB) media plate were harvested and resuspended in sterile distilled water to a final concentration of OD_600_=0.02. The bacterial suspension was dispensed into plates containing *A. thaliana* seedlings, and plates were incubated for 3 min before bacteria were removed by decantation.The entire rosette of each inoculated seedling was collected 3 days post-inoculation (dpi), and their fresh weight was measured before surface-sterilization (5% H_2_O_2_ for 3 min, followed by 3 washes with sterile distilled water) for colony assays. For syringe-inoculation, *Pst* DC3000 *avrRpt2* or *Pst* DC3000 *avrRps4* were harvested from KB plates and resuspended in 10 mM MgCl_2_ to a final concentration of OD_600_=0.0005. The suspension was infiltrated into leaves of 4-week-old seedlings with a needleless syringe. Bacterial growth was determined at 3dpi by colony counting.

### Confocal microscopy

Five-day-old *A. thaliana* seedlings were imaged on a TCS SP8 confocal microscope (Leica, Wetzlar, Germany) with a 40x water corrected objective (1.10 NA) and a 488 nm excitation laser at 5% intensity. GFP emission was captured from 499 to 559 nm with a photomultiplier tube, at a gain of 450.3. Propidium Iodide emission was captured from 576 to 691 nm with a Hybrid detector, at a gain of 55.3. For the BFA assay, the laser intensity was reduced to 2%. GFP emission was captured from 507 to 539 nm with a Hybrid detector, at a gain of 10. FM4-64 emission was captured from 705 to 755 nm with a Hybrid detector, at a gain of 50. Images were combined to a frame average of 4. HEK293 cells transfected with plasmid DNA (pcDNA3.2, CMV::GFP, CMV:: opti2-ACD6^Col-0^-mGFP5-6xHis, or CMV::MHAL-mGFP5-6xHis) and incubated for 24 h were visualized through Carl Zeiss 880 Confocal microscopy system (Carl-zeiss, Oberkochen, Germany) at 20X magnification and a 488 nm excitation laser at 5% intensity. GFP emission was captured from 490 to 543 nm with a photomultiplier tube at a gain of 500 (for GFP) or 1000 (for ACD6^Col-0^-mGFP5-6xHis and MHAL-mGFP5- 6xHis).

### Subcellular fractionation

We followed a published protocol^81^ with modification. About 100 mg of 10-d-old *A. thaliana* seedlings were ground on ice with pestle and mortar. 200 µL sucrose buffer (20 mM Tris [pH 8], 0.33 M sucrose, 1 mM EDTA, 1 mM DTT and protease inhibitor cocktail) was added. Samples were spun at 2,000 g for 10 min at 4°C to remove plant debris. 45 µl supernatant was removed as total lysate fraction (T). The rest was centrifuged at 20,000 g for 1 h at 4°C. 45 µl of the resulting supernatant was designated as soluble fraction (S). The pellet was resuspended in 45 µl sucrose buffer and considered as microsomal fraction (M).

### Split-luciferase complementation assay

We followed a published protoco^58^. *Agrobacterium tumefaciens* GV3101 with expression constructs for ACD6- cLuc, ANK-cLuc, BAK1-cLuc, MHA1-Est-1-nLuc, MHA1-Ty-0-nLuc or MHA1L-nLuc were suspended at 5 × 10^8^ cfu/ml in ½ MS medium. The suspension was infiltrated into the leaves of 30-d-old *Nicotiana benthamiana* plants. For CCD imaging, leaves were sprayed 48 h later with 1 mM D-Luciferin-K (PJK GmbH, Kleinblittersdorf, Germany) in water containing 0.02% Silwet L-77 (Helena) and kept in the dark for 10 min before measurement. The luminescence signal was recorded with an Orca 2-BT cooled CCD camera (Hamamatsu Photonics, Shizuoka, Japan). The experiment was repeated twice.

### Co-immunoprecipitation

ACD6-FLAG was co-expressed with GFP-MHA1 and GFP-MHA1L in protoplasts isolated from 4-week-old *Arabidopsis thaliana* transgenic plants. Total protein was extracted with extraction buffer (50 mM HEPES [pH 7.5], 150 mM KCl, 1 mM EDTA, 0.5% Triton-X 100, 1 mM DTT, and proteinase inhibitor cocktail). For anti-GFP co-IP, the extracted proteins were incubated with GFP-Trap_M (ChromoTek, Planegg, Germany) at 4°C under gentle rotation for 2 h. The magnetic particles were washed seven times with washing buffer (50 mM HEPES [pH 7.5], 150 mM KCl, 1 mM EDTA, 0.5% Triton-X 100, 1 mM DTT) at 4°C. Proteins were detected by immunoblotting using anti-GFP (Santa Cruz, California, USA) and anti-FLAG (Sigma-Aldrich, MO, USA) antibodies.

### In vitro pull-down assays

GST-, MBP-6xHis-, and 6xHis-fusion proteins were expressed in *E. coli* BL21. Bacterial cultures in LB were incubated at 28°C until they had reached OD_600_=0.6, when protein expression was induced with 0.4 mM IPTG at 20°C for 16 hr. Tagged proteins were purified separately using glutathione agarose beads (GE Healthcare, Chicago, USA) or Ni-NTA affinity agarose beads (QIAGEN). Purified proteins were dissolved in a buffer containing 25 mM Tris-HCl (pH 7.5), 100 mM NaCl, and 1 mM DTT and concentrated with Amicon Ultra-15 Centrifugal Filter Units (Millipore, Darmstadt, Germany). For GST pull-down 2 µg GST-tagged Protein, 20 µl Glutathione Sepharose 4B (GE Healthcare) and 10 µg MBP-6xHis-tagged protein were taken up in 1 ml pull-down buffer (25 mM Tris-HCl [pH 7.5], 200 mM NaCl, 0.5% [v/v] Triton X-100) and incubated for 4 h under gentle rotation. For washing, the beads with bound protein were spun down (500 rpm, 10 min), the supernatant was discarded, beads were resuspended in pull-down buffer and incubated for 1 h under gentle rotation. Washing was repeated 6 times. Elution of GST-tagged protein from beads was performed by resuspending pelleted beads (500 rpm, 10 min) in 100 µl Elution buffer (25 mM Tris-HCl [pH 7.5], 150 mM NaCl, 30 mM glutathione) and incubated for 1 h under gentle rotation. After centrifugation (1,000 rpm, 20 min), GST-tagged protein eluate was collected from supernatant. For MBP pull-down, 2 µg MBP-tagged protein, 20 µl Amylose resin (New England Biolabs), 5 µg GST-tagged protein, and 5 µg 6xHis-tagged protein were taken up in 1 ml pull-down buffer with the same incubation and washing method, and eluted with 100 µl Elution buffer (25 mM Tris-HCl [pH 7.5], 150 mM NaCl, 50 mM maltose).Protein purification and pull-down were performed at 4°C. Proteins were detected by immunoblotting using anti-Glutathione-S-Transferase (Sigma-Aldrich), anti-polyHistidine-Peroxidase (Sigma-Aldrich) and anti-MBP (AbClonal, Wuhan, China) antibodies.

### Blue Native-PAGE

Blue native polyacrylamide gel electrophoresis (BN-PAGE) was performed using the Bis-Tris NativePAGE system (Invitrogen, CA, USA) according to manufacturer’s instructions. Briefly, eight 12-d-old seedlings were collected and ground in 1×NativePAGE Sample Buffer (Invitrogen) containing 0.5% n-dodecyl β-D-maltoside (DDM) and protease inhibitor cocktail, followed by 13,000 rpm centrifugation for 20 min at 4°C. 15 µl supernatant mixed with 0.3 µl 5% G-250 Sample Additive was loaded and run on a NativePAGE 3-12% Bis-Tris gel. Native gels were transferred to PVDF membranes (Millipore, Darmstadt, Germany) using NuPAGE Transfer Buffer, followed by protein blotting. For the second dimension of electrophoresis, a 5.7 cm strip of BN-PAGE gel was incubated in Laemmli sample buffer (50 mM Tris-HCl [pH 6.8], 100 mM DTT, 2% (w/v) SDS, 0.1% bromophenol blue, 10% (v/v) glycerol) for 10 min, microwaved for 20 s, and then rotated for another 5 min before loading the strip into the well of a NuPAGE 4-12% Bis-Tris protein gel (Invitrogen).

### Protein homology search and alignment

HHPred search and structure modeling with MODELLER were performed online (https://toolkit.tuebingen.mpg.de/#/tools/hhpred), using the MPI Bioinformatics Toolkit^30^. HHPred default search parameters were used for searches with ACD6-Col-0 (excluding residues 1-90) as query. Alignments were visualized with Jalview^63^. Default parameters were used for homology modeling with MODELLER^82^. We used the SMART database (http://smart.embl-heidelberg.de/) to predict the boundaries of the ankyrin repeats in ACD6, and the TMHMM server v2.0 (http://www.cbs.dtu.dk/services/TMHMM/) to predict transmembrane domains. Structures were visualized using PyMOL^66^.

### Injection of oocytes and electrophysiological measurements

Dissected and preselected *Xenopus laevis* oocytes were obtained from Ecocyte Bioscience (Dortmund, Germany). Oocytes were kept in ND96 (96 mM NaCl, 2 mM KCl, 1 mM MgCl_2_, 1.8 mM CaCl_2_, 2.5 mM sodium pyruvate, 5 mM HEPES adjusted to pH of 7.4 with NaOH). cRNA was produced from *Mlu* I linearized and phenol-chloroform purified pOO2 plasmids ^60^ containing the coding sequences of ACD6-1, ACD6-Col-0 or MHA1L using the mMESSAGE mMACHINE SP6 Transcription Kit (Life Technologies GmbH, Darmstadt; Germany) following manufacturer’s instructions. Oocytes were injected with 50 nl cRNA with a concentration of 300 ng/µl for ACD6 variants and 0.1 ng/µl for MHA1L. Oocytes were kept in ND96 for 3-4 d at 18°C. Electrophysiology was performed in a small recording chamber containing the recording solution.

For sodium transport measurements the test solution contained 110 mM choline chloride, 2 mM CaCl_2_, 2 mM MgCl_2_, 5 mM N-morpholinoethane sulfonate (MES), pH adjusted to 5.5 with Tris(hydroxymethyl) aminomethane (TRIS) with and without 10 mM NaCl. Recording solutions for calcium-induced chloride currents were: -Cl solution (100 mM sodium glutamate, 5 mM MES, pH adjusted to 5.5 with TRIS); -Cl/+Ca solution (90 mM sodium glutamate, 10 mM CaNO_3_, 5 mM MES, pH adjusted to 5.5 with TRIS); -Ca solution (100 mM sodium chloride, 5 mM MES, pH adjusted to 5.5 with TRIS) and +Ca solution (90 mM sodium chloride, 10 mM CaCl_2_, 5 mM MES, pH adjusted to 5.5 with TRIS).

Two-electrode voltage-clamp measurements were performed using 3 M KCl-filled glass capillaries of around 2 Mohm resistance of the electrodes. For measurements in -Cl solutions, an agarose bridge was used to connect the recording chamber with a separated reference electrode bathed in 3 M KCl solution via a glass capillary filled with 3 M KCl and 2 % agarose. Currents were recorded with 100 ms steps over a range from +80 mV to -140 mV in 10 or 20 mV decrements. Electrical recordings were performed after complete wash of the bathing solutions to test solutions and the mean induced currents by the test solutions are shown (background currents were subtracted). Data from representative oocyte batches are shown.

### Fura-2 imaging of HEK293 cells

The constructs *CMV::ACD6*^Col-0^ and *CMV::MHA1L* were synthesized in pcDNA3.1 myc-6xHis or pcDNA3.1 mGFP5-6xHis (GenScript Biotech, NJ, USA). Two codon-optimized ACD6^Col-0^ sequences were obtained and synthesized from gBlock synthesis (GenScript) and subcloned into pcDNA3.1 myc-6xHis or mGFP5-6xHis through the HiFi assembly method (NEB). The Q5 Site-directed mutagenesis kit (NEB) was used to generate *CMV::opti2-ACD6-1* and *CMV::opti2-ACD6^L557F^*. The constructs were verified through whole plasmid sequencing (Plasmidsaurus, OR, USA). Transfection experiments used plasmid DNA extracted from *E. coli* strain Stbl3 (Thermo Fisher Scientific) using the Pureyield Plasmid Miniprep system (Promega).

Fura-2 signals in human embryonic kidney 293 (HEK293) cells were studied as described^23, 40–42^. HEK293 cells were obtained from Duke Cell Culture Facility and maintained in DMEM medium supplemented with 10% fetal bovine serum, 1% penicillin and streptomycin in a cell culture incubator (Thermo Fisher Scientific) at 37°C and 5% CO_2_. HEK293 cells were seeded and grown on poly-lysine-coated Lab-Tek II eight-well chamber slides (NUNC) overnight and transfected with plasmid DNA using the Lipofectamine™ 3000 Transfection System (Invitrogen) according to manufacturer’s instructions. Cells were loaded with Fura-2AM (5 mM; Sigma Aldrich), and Fura-2 signals were imaged 22-26 h after transfection using an inverted fluorescence microscope (Axiovert 200; Zeiss) with 20X magnification, equipped with two filter wheels (Lambda 10-2; Sutter Instruments, CA, USA), and a cooled CCD/CMOS camera (CoolSNAPfx/Prime 95B; Teledyne Photometrics, CA, USA)^23, 42^. Ratiometric emission images at 510 nm with excitation of 340 nm and 380 nm, respectively, were collected using MetaFluor software^23, 42^ or Micro-Manager software (https://micro-manager.org/)^83^. The ratio between F340:F380 was calculated from 50-70 cells selected based on highest apparent signal increases using MetaFluor or Image J. After Fura-2AM loading into cells, the cells were incubated in standard buffer (130 mM NaCl, 3 mM KCl, 0.6 mM MgCl_2_, 10 mM glucose, 10 mM HEPES pH7.4 [adjusted with NaOH]), and 0.1 mM Ca^2+^ for 30 minutes. After imaging for 25 s, the concentration of Ca^2+^ was increased to 2.5 mM by adding a standard buffer supplemented with Ca^2+^ into the chamber. Additional images were collected for the next 305 s.

### Calcium influx assays

For aequorin luminescence measurements, an aequorin transgene^84^ was introduced into the *acd6*-1*, mha1l, p35S::MHA1L,* and *dnd1-*1 backgrounds by crossing, and homozygous F_2_ progeny identified. *acd6*-ko (Col-0) lines in *p35S::aequorin* background was generated by CRISPR/Cas9 mentioned above. Est-1 and *acd6*-ko (Est-1) were introduced with a *pUBI10::YFP-aequorin* expression vector^85^, which carry a hygromycin seletion marker. The genotyping primers for the aequorin transgene insertion, *acd6*-ko, and *acd6*-1 are listed in **Table S2**. The *p35S::MHA1L* transgene carries the fluorescent co-dominant marker FAST (fluorescence-accumulating seed technology)^59^. Individual surface-sterilized seeds were sown directly into 96-well white plates containing 50 µl ½ MS in each well. The media was changed to 50 µl of 10 μM coelenterazine (Yeasen, 40904ES02) in sterile water 5 day after germination. Seedlings were kept in the dark at room temperature for 16 h before treatment. The background luminescence of seedling was recorded for 2 min before application of flg22 and chitin (5 s for NaCl) on a microplate reader (BioTek Synergy H1). Then 50 μl of 200 nM flg22, 200 μg/ml chitin, or 600 nM NaCl was applied to each well using an auto-syringe injection system, and luminescence was recorded for a further 20 min in 25 s intervals (20 s in 0.5 s interval for NaCl treatment). The remaining aequorin of the sample was discharged with 100 μl 2 M CaCl2 (in 20% ethanol). The treatment-induced luminescence counts relative to the remaining total luminescence counts (L/L_Max_) were used to estimate calcium influx. For pathogen treatment, bacteria were grown to OD_600_=3.0 in KB medium, pelleted at 4,000 rpm and resuspended in 10 μM coelenterazine. The suspension was infiltrated into leaves of 4-week-old *A. thaliana* plants with a needleless syringe. Leaf punches (6 mm in diameter) from the infiltrated leaves were incubated with 150 μl 10 μM coelenterazine in a 96-well white plate, and luminescence was recorded for 6 h in a 2 min interval. The remaining aequorin was discharged with 150 μl 2 M CaCl_2_ (in 20% ethanol).

For R-GECO1(ref. ^46^) luminescence measurement, cotyledons from 10-day-old seedlings were removed 12 h prior to imaging and placed in sterile deionized water in a Hybriwell passive microfluidic device (Grace Bio-Lab, OR, USA) fixed to a 24 x 50mm No. 1.5 coverslip. Samples were imaged using a Zeiss 880 Airyscan Inverted confocal on a Zeiss Axio Observer Z1 microscope using a Zeiss EC Plan-Neofluar 10x/0.30 M27 objective. Times series images were collected using 458 nM (for mTurquoise2) and 561 nm (for R-GECO1) laser lines for excitation, and the Zeiss 880 S1 channel set to 463-544 nm (for mTurquoise2) and the S2 set to 588-651 nm (for R-GECO1). The pinhole was set to 8.43 airy units for each channel, line averaging of 4, 512 x 512 pixel image format, and 0.6x optical zoom. Time series was acquired with 3.77 scan time for each image at 5 second intervals. At t = 0, 60 µl of 100 nM flg22 was added to the cotyledons. Ratiometric image processing was performed using a custom ImageJ macro^86^. Briefly, the macro performs background subtraction and segmentation, then automatically divides the R-GECO1 channel by the mTurquoise2 channel to obtain a ratio image. Normalized RGECO1/mTurquoise2 values were obtained by obtaining the average signal prior to flg22 treatment (R_0_) and using the following formula (R-R_0_)/R_0_, where R is the ratio value at each time point, to determine the normalized signal.

### DATA AND SOFTWARE AVAILABILITY

DNA sequences are available at Genbank: *ACD6*-Ty-0 MH120293, *MHA1-*Ty-0 MH120291, *MHA1*-Est-1 MH120292.

## ACKNOWLEDGEMENTS

We thank past and present members of the Weigel lab, Katherine Huffer and Kenton Swartz, and Yalong Guo for discussion, and Alejandra Duque-Jaramillo, Farid El-Kasmi, Derek Lundberg, Thorsten Nürnberger, Wei Yuan and Jingbo Zhang for comments on the manuscript. We thank Maricris Zaidem for Ty-0 x Est-1 F_2_ seeds, Polina Novikova for *Arabidopsis lyrata* resequencing data, Xin Li for helper-NLR mutant seeds, Yan Guo for *pUBI::YFP-aequorin* plasmid, and Dongqin Chen for the aequorin transgenic line. We thank Monika Demar, Frederik Unger, Sonja Kersten, Kai Wang and Martin Bayer for help with experiments, and the ZMBP Analytics Laboratory at the University of Tübingen for SA quantification. This work was supported by National Key Research and Development Program, Ministry of Science and Technology of China (No 2022YFD1201802), the Ministry of Education of China (the 111 Project B13006) and the 2115 Talent Development Program of China Agricultural University (No 2020RC013) (W.Z.), an EMBO Long-term Fellowship (969-2016 to L.L.), an HFSP Long-term Fellowship (LT000314/2017-L to L.L.), USDA NIFA Postdoctoral Fellowship Award Number 2021-67012-35140 (R.H.), the DFG (NE 1727/2-2 to B.N.), the Ministry of Science, Research and Arts Baden-Württemberg through the Regio Research Alliance “Yield Stability in Dynamic Environments” (U.Lud., D.W.), the DFG through SFB1101, and the Max Planck Society (D.W.).

## AUTHOR CONTRIBUTIONS

Conceptualization: J.C., L.L., J.K., U.Lud., S.H., Z.P., D.W., W.Z. Methodology: J.C., L.L., J.K., B.N. R.H., W.Z. Formal analysis: W.Z. Investigation: J.C., L.L., J.K., B.N., M.W., Y.C., W.Z. R.H., M.T., L.W., K.V., M.E.A., C.L., J.K., A.C.B, R.S., U. Lutz. Writing–original draft: W.Z., D.W. Writing–review, and editing: W.Z., J.C., L.L., U.Lud., D.W. Supervision: S.H., Z.P., U.Lud., D.W., W.Z. Project administration: D.W., W.Z. Funding acquisition: W.Z., L.L., B.N., U.Lud., D.W.

## SUPPLEMENTAL FIGURES

**Figure S1.**
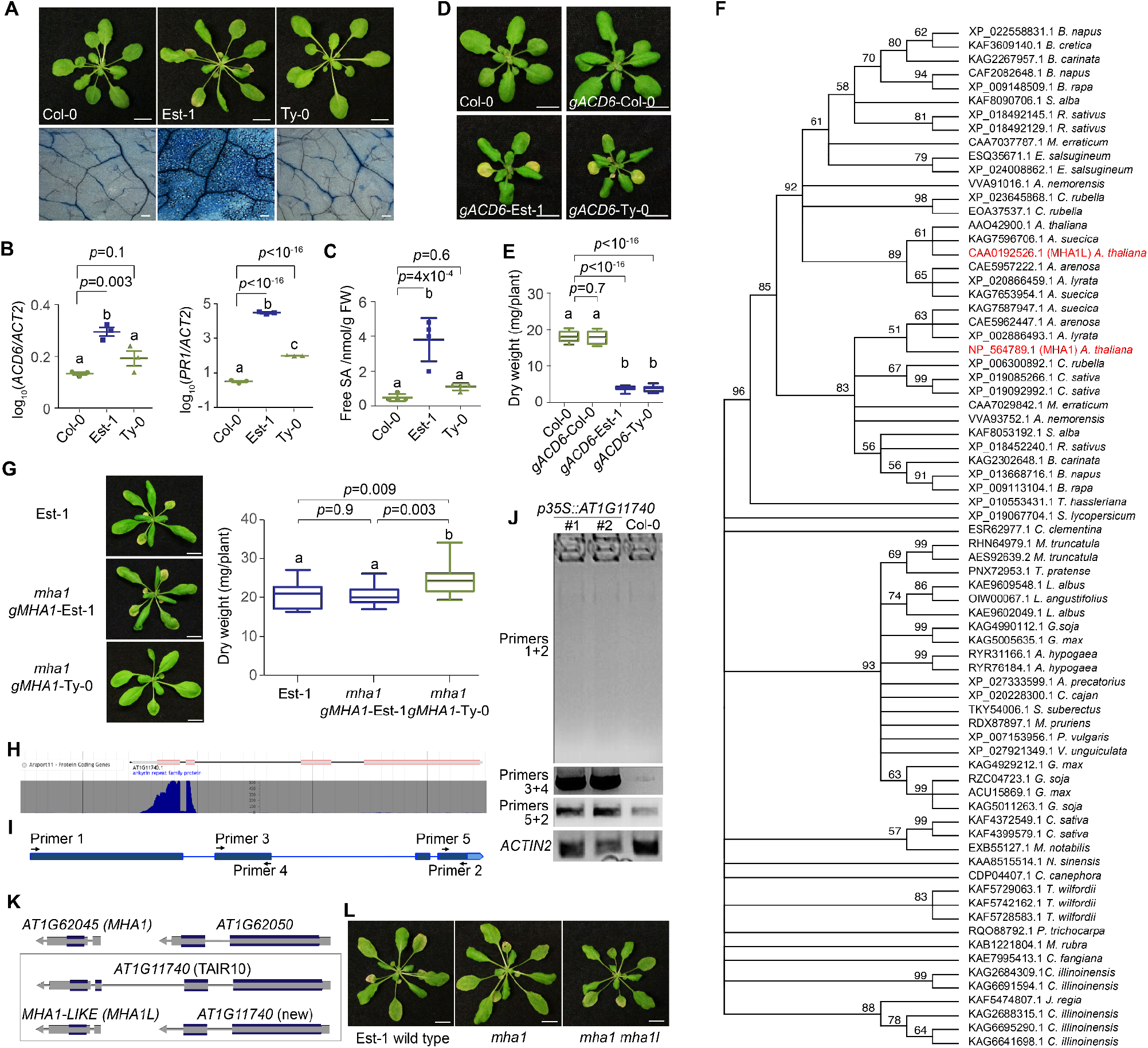
Identification of *MHA1* as a genetic modifier of *ACD6-*Est-1, related to Figure 1. **(A)** Top, four-week-old plants with different *ACD6* alleles. Bottom, close-up of the sixth leaf, stained with Trypan blue for dead cells. Scale bars 1 cm (top), 0.2 mm (bottom). **(B)** *ACD6* and *PR1* expression in different genotypes, as measured by qRT-PCR from three biological replicates each. **(C)** Accumulation of free salicylic acid (SA) in different genotypes. **(D)** Representative 18-day old transgenic plants expressing different genotypes of genomic *ACD6* fragments in Col-0 background in 23°C LD. Scale bars 1 cm. **(E)** Dry weight of 3-week-old plants from (D). **(F)** Phylogeny of 74 MHAI-like proteins from dicots and monocots. MHA1 and MHA1L from *A. thaliana* are in red. **(G)** Transgenic Est-1 *mha1* mutants containing genomic *MHA1* fragments from Est-1 or Ty-0. Left: Plant phenotypes. Scale bar, 1 cm. Right, Leaf biomass of 4-weekold plants. **(H)** RNA-seq data from light-grown seedlings indicate very different coverages of the 5’ and 3’ ends of the annotated *AT1G11740* gene model. **(I, J)** RT-PCR analysis of Col-0 transformant overexpressing the annotated *AT1G11740* gene model. Full-length *AT1G11740* cDNA (primers 1+2) cannot be amplified from T_2_ transgenic plants or Col-0 plants. A 5’ product (primers 3+4) over accumulates in T_2_ transgenic plants, while a 3’ product (primers 5+2) is increased much less in transgenic compared to Col-0 non-transgenic plants. **(K)** Correction of gene annotation of *MHA1- LIKE* (*MHA1L*) in the *A. thaliana* reference genome. Paralogous *MHA1* region shown on top. **(L)** CRISPR/Cas9 *mha1* and *mha1/mha1l* mutants in Est-1 background. *p*-values from Tukey’s HSD test (B, C, E, G). Letters indicate significantly different groups, as determined by one-way ANOVA followed by Tukey’s test at α = 0.05 (B, C, E, G).

**Figure S2.**
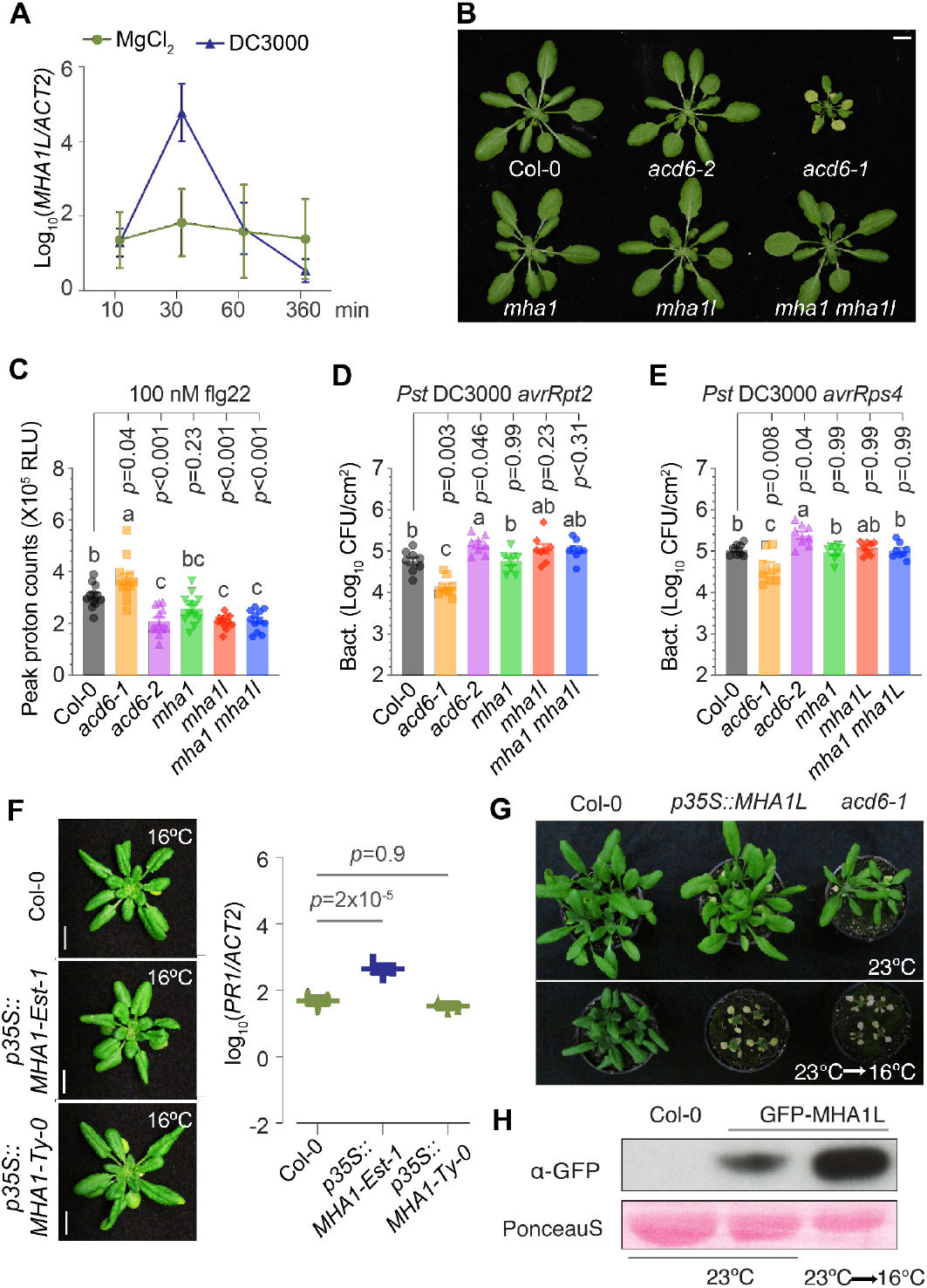
MHA1s were required for PTI and basal resistance, related to Figure 2. **(A)** Time course of *MHA1L* mRNA accumulation, as measured by qRT-PCR, in Col-0 plants after infiltration with *Pst* DC3000 (OD_600_=0.3). **(B)** Representative plant morphology of different genotypes. Plants were grown at 23°C. **(C)** Peak value of flg22-induced ROS production in Figure 2A. **(D, E)** Bacterial growth of *Pst* DC3000 *avrRpt2* (D), and *Pst* DC3000 *avrRPS4* (E) at 3 days after inoculation in indicated plants. Data are represented as mean ± s.e.m.; n = 8. **(D)** Plants overexpressing *MHA1*-Est-1 or *MHA1*-Ty-0 alleles in the Col-0 reference background grown at 23°C. Left, plant phenotypes, Right, accumulation of *PR1* mRNA, as measured by qRT-PCR from three biological replicates each. **(E)** Temperature-dependent phenotypes of *p35S::GFP-MHA1L* and *acd6*-1 plants. Top, 3-week-old plants grown in 23°C long days. Bottom, plants grown for 10 days in 23°C long days and then moved to 16°C for 10 days. **(F)** GFP-MHA1L protein accumulation in 2-week-old plants in Col-0 background grown in 23°C long days or plants grown in 23°C long days for 12 days, then moved to 16°C for 2 days. *p*-values from Tukey’s HSD test (C, D, E, F). Letters indicate significantly different groups, as determined by one-way ANOVA followed by Tukey’s test at α = 0.05 (C, D, E).

**Figure S3.**
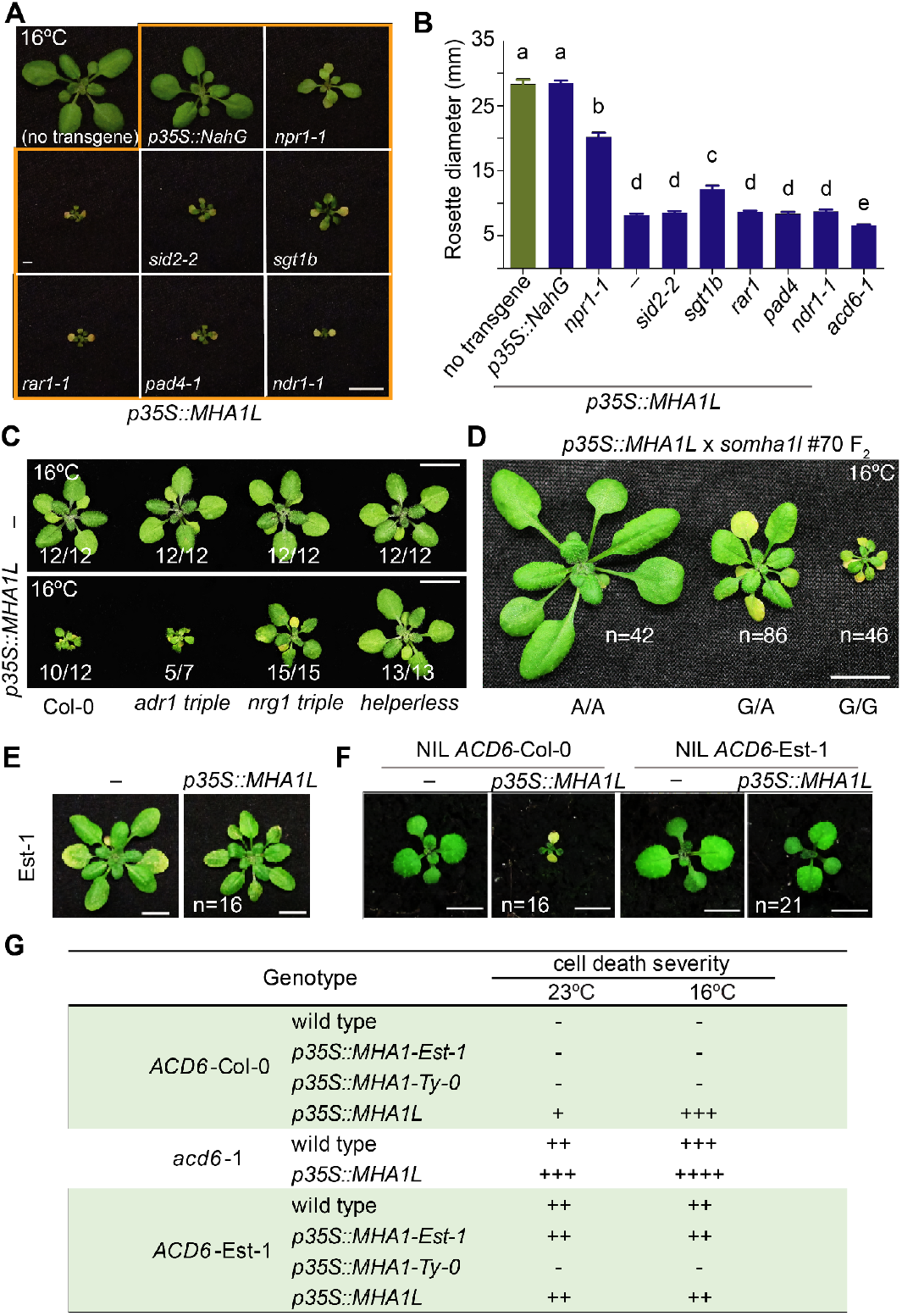
Identification of Suppressor of MHA1L, related to Figure 3. **(A)** Representative *p35S::MHA1L* T_2_ transformants in different backgrounds (thick orange boundary), with non-transgenic Col-0 as control on the top left. Plants were grown at 16°C. **(B)** Rosette diameter of T_2_ transformants. At least 15 plants were used for each measurement. *acd6*-1 for comparison. Letters above data points indicate significantly different groups, as determined by one-way ANOVA followed by Tukey’s test at α = 0.05. **(C)** Effects of mutating *hNLR* genes in Col-0 or *MHA1L* overexpressors. Numbers indicate fractions of T_1_ transgenic plants with the phenotype shown. Scale bars 1 cm. **(D)** Co-segregation between the polymorphism at codon 575 of *ACD6* with *p35S::MHA1L*-associated cell death in a backcross F_2_ population derived from *somha1l* suppressor mutant #70 and the parental *p35S::MHA1L* line. “A” corresponds to the *ACD6* wild-type allele.**(E)** Four-week-old *p35S::MHA1L* T_1_ transformants grown under 16°C long days are shown next to non-transgenic controls. The phenotypic effects of *p35S::MHA1L* are suppressed in an Est-1 background. Compare to Figure 2E. **(F)** The phenotypic effects of *p35S::MHA1L* are apparent in a NIL^1^ with the *ACD6*-Col-0 allele, but not in a NIL with the *ACD6*-Est-1 allele. **(G)** Summary of genetic interactions between *MHA1*, *MHA1L* and *ACD6*.

**Figure S4.**
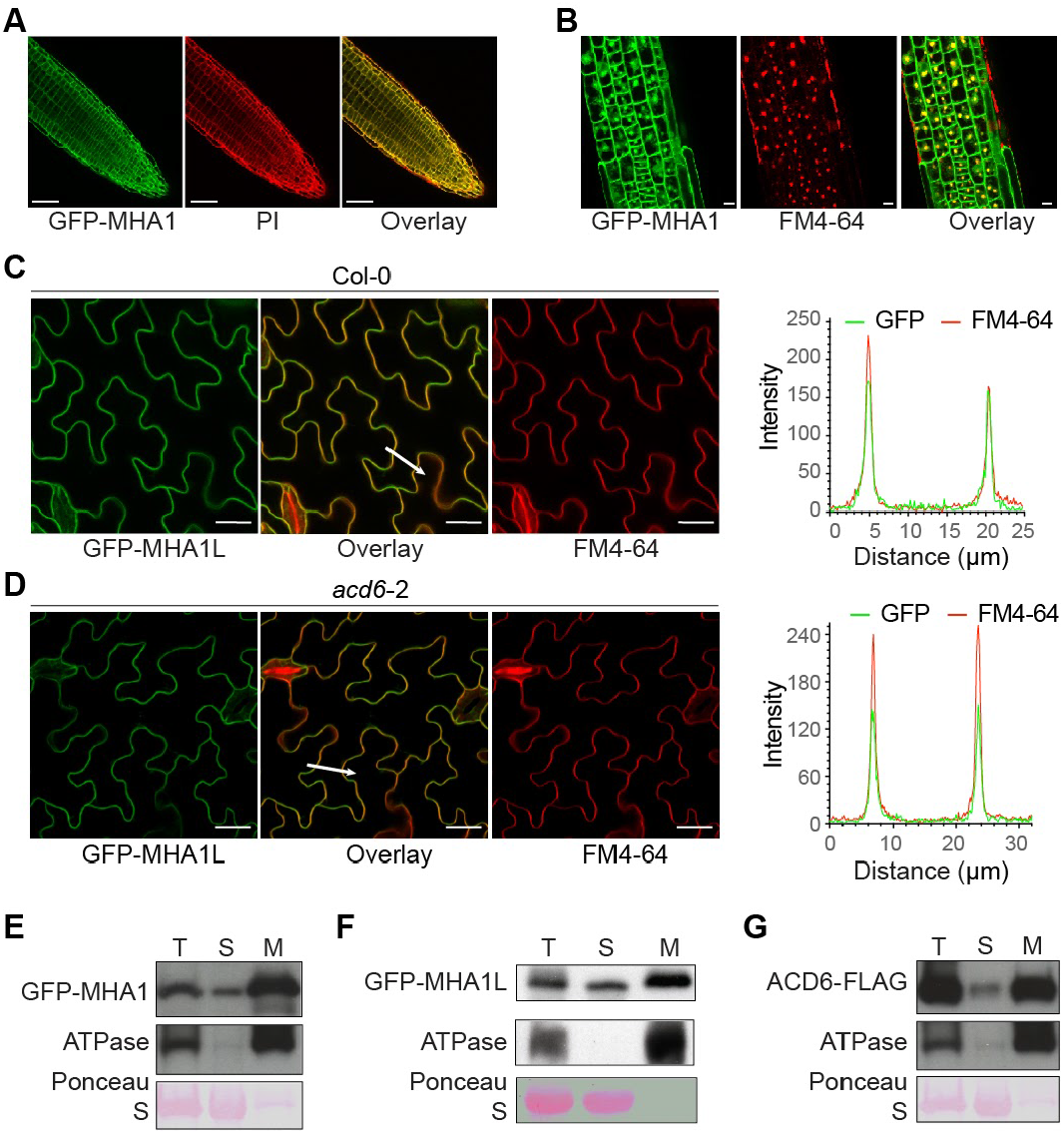
MHA1 and MHA1L localise to both cytoplasm and plasma membranes, related to Figure 4. **(A)** Localization of GFP-MHA1-Est-1 in root cells; cell walls were stained with propidium iodide (PI). Scale bars 50 µm. **(B)** Change of subcellular localization of GFP-MHA1-Est-1 in response to 10 µM BFA (45 min after treatment). Plasma membranes were stained with FM4-64. Scale bars 5 µm. **(C-D)** Confocal images of FM4-64-stained cotyledon epidermal cells of GFP-MHA1L transgenic plants. The white arrows indicate the lines for the fluorescence intensity profiles, which are shown on the right. Scales, 20 µm. **(E-G)** Subcellular microsomal fractionation of extracts from *p35S::GFP-MHA1* (E), *p35S::GFP-MHA1L* (F) and *pACD6::ACD6-*Col-0*-FLAG* (G) plants. T, total; S, supernatant; M, microsomal fraction.

**Figure S5.**
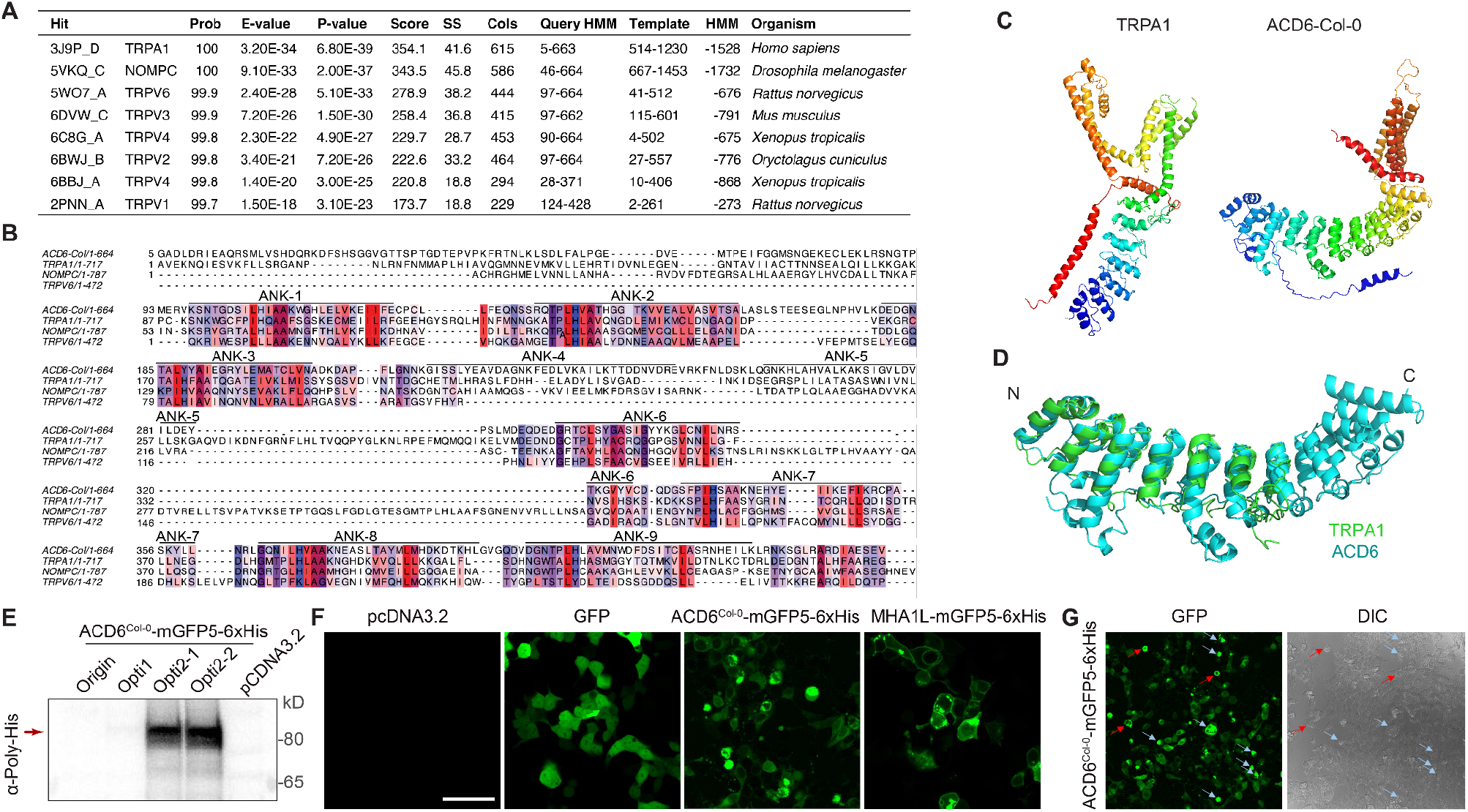
Homology modeling and visualization of ACD6, related to Figure 5. **(A)** HHPred hits with ACD6 from Col-0 as a query. **(B)** Protein sequence alignment of ankyrin (ANK) repeats of ACD6, fly NOMPC, human TRPA1 and rat TRPV6. Coloring indicates hydropathy index. **(C)** Structure of TRPA1 (3J9P) compared to homology-modeled ACD6 (Q8LPS2) by AlphaFold 2 ^87^. . Structures are colored from N to C terminus (blue to red). Note that the C-terminal portion of the ACD6 model does not model well on the TRPA1 pore. **(D)** Superimposed structures for ANK domain of TRPA1 and ACD6. **(E)** Validation of codon-optimized ACD6^Col-0^-mGFP5-6xHis in HEK293 cells transfected with pcDNA3.2 empty vector, ACD6^Col-0^-mGFP5-6xHis or codon-optimized ACD6^Col-0^- mGFP5-6xHis constructs (opti1 and opti2) through immunoblot. The arrow indicates ACD6^Col-0^-mGFP5-6xHis protein. **(F)** Confocal microscopy to visualize ACD6^Col-0^-mGFP5-6xHis and MHA1L-mGFP5-6xHis proteins through confocal microscopy in HEK293 cells transfected with pcDNA3.2 empty vector, or expressing GFP, codon-optimized ACD6^Col-0^-mGFP5-6xHis, or MHA1L- mGFP5-6xHis. Scale bars 50 µm. **(G)** Correlation between strong ACD6^Col-0^-GFP signal and dying cells. Dying cells tended to have higher expression levels of ACD6-mGFP5 (light blue arrows), and the localization pattern of ACD6-mGFP5 tended to be more membrane-enriched in dying cells (red arrows) than in low expressing cells.

**Figure S6.**
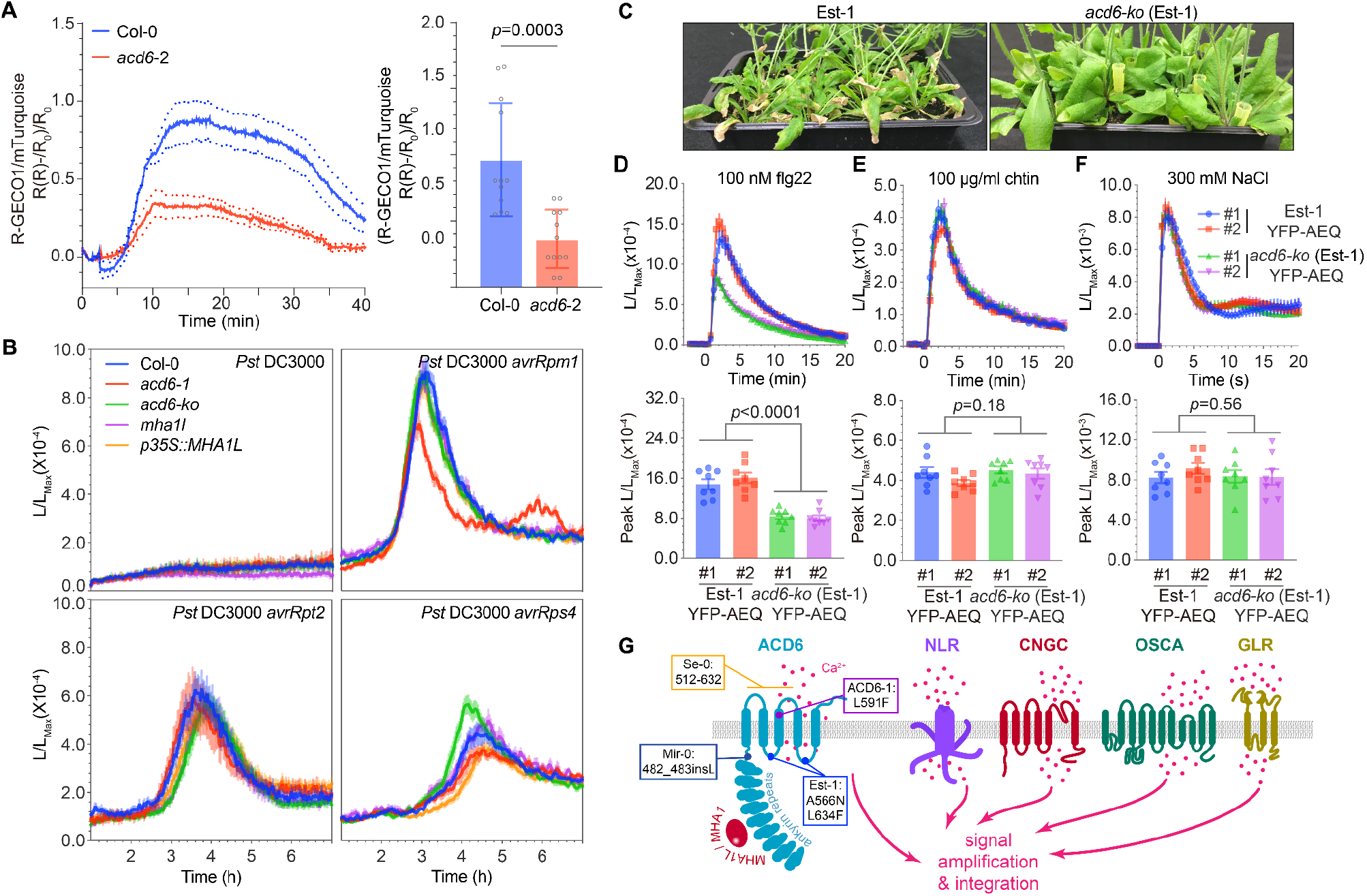
Requirements of ACD6 and MHA1L for PAMP- and effector-induced calcium signaling, related to Figure 6. **(A)** Cotyledons of 10-day old seedlings were imaged. 100 nM flg22 was applied at t=0 min and normalized R-GECO1/mTurquoise2 (R-R_0_)/R_0_ signal was quantified (n=12). Maximum R-GECO1/mTurquoise2 peak values that were obtained for individual repeats (n=12, *p*-values from Student’s t-test). Experiments were repeated twice, with similar results. **(B)** Time course of aequorin-luminescence after bacteria infiltration (OD_600_=0.1). Data are means ± s.e.m. (n≥8). **(C)** Representative morphology of wild-type Est-1 and Est-1 *acd6-ko* plants. **(D, E, F)** Top, time course of aequorin-luminescence after 100 nM flg22 (D), 100 μg/ml chitin (E), or 300 mM NaCl (F) treatment, which were applied at t= 0. Bottom, peak values of each sample. Data are means ± s.e.m. (n=8). *p*-values from Student’s t-test. **(G)** Biochemical model for ACD6 function. ACD6 facilitates calcium influx, either by regulating ion channels or acting as an ion channel itself, and its activity is enhanced either by binding of MHA1L to its ankyrin repeats, or by mutations in the transmembrane domains. MHA1-Ty-0 can reduce activity of the ACD6-Est-1 variant by binding to the ankyrin repeats. Locations of changes in induced and natural gain-of-function alleles are indicated. Other classes of calcium channels regulating immunity are shown as well; the extent of redundancy and cross-regulation between them is an open question.

## SUPPLEMENTAL TABLES

**Table S1.**
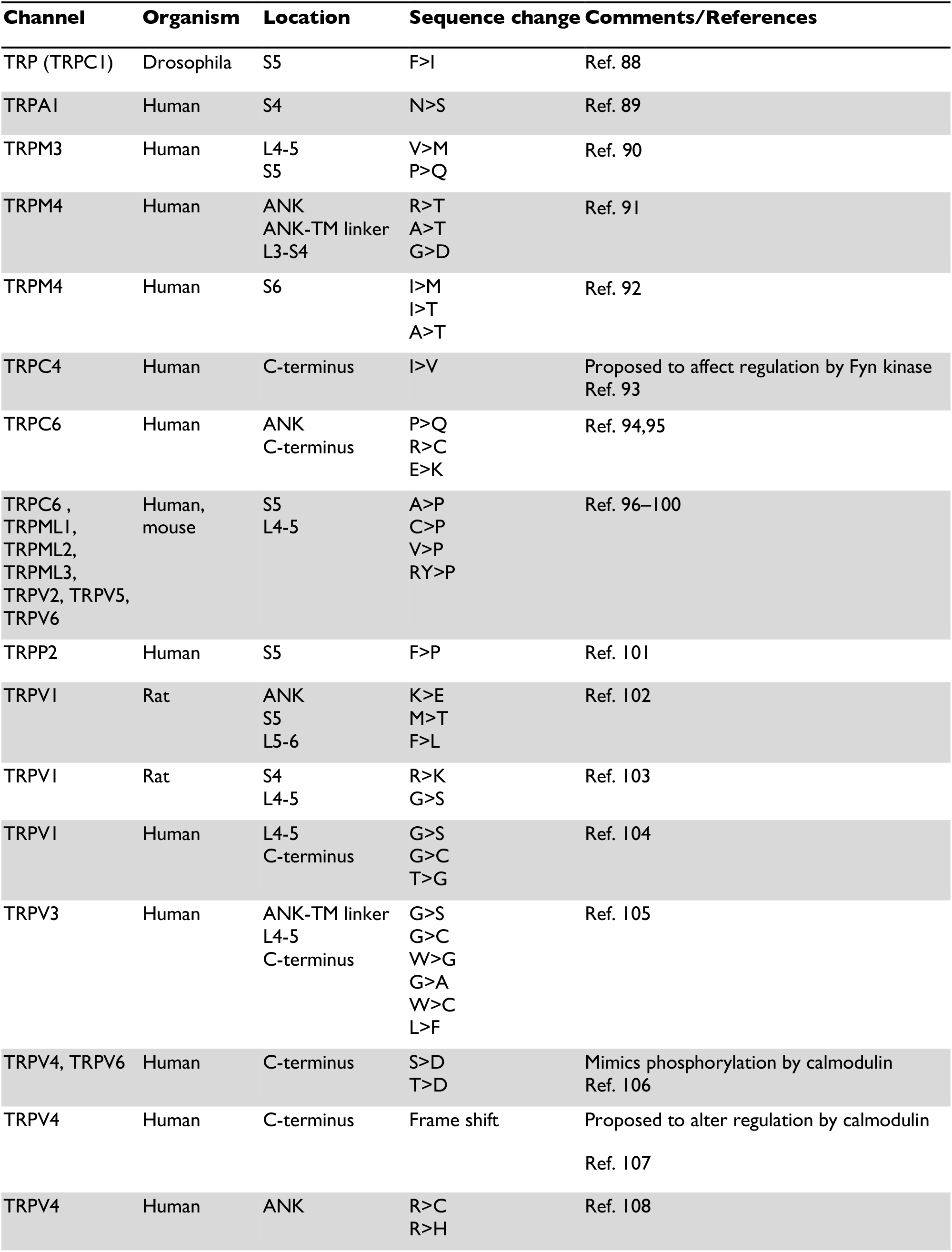
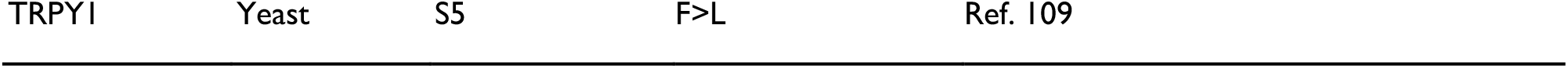
Gain-of-function mutations in TRP channel genes, related to Figure 3 and 5.

| Channel | Organism | Location | Sequence change | Comments/References |
| --- | --- | --- | --- | --- |
| TRP (TRPC1) | Drosophila | S5 | F>I | Ref. 88 |
| TRPA1 | Human | S4 | N>S | Ref. 89 |
| TRPM3 | Human | L4-5<br>S5 | V>M<br>P>Q | Ref. 90 |
| TRPM4 | Human | ANK<br>ANK-TM linker<br>L3-S4 | R>T<br>A>T<br>G>D | Ref. 91 |
| TRPM4 | Human | S6 | I>M<br>I>T<br>A>T | Ref. 92 |
| TRPC4 | Human | C-terminus | I>V | Proposed to affect regulation by Fyn kinase<br>Ref. 93 |
| TRPC6 | Human | ANK<br>C-terminus | P>Q<br>R>C<br>E>K | Ref. 94,95 |
| TRPC6 ,<br>TRPML1,<br>TRPML2,<br>TRPML3,<br>TRPV2, TRPV5,<br>TRPV6 | Human,<br>mouse | S5<br>L4-5 | A>P<br>C>P<br>V>P<br>RY>P | Ref. 96–100 |
| TRPP2 | Human | S5 | F>P | Ref. 101 |
| TRPV1 | Rat | ANK<br>S5<br>L5-6 | K>E<br>M>T<br>F>L | Ref. 102 |
| TRPV1 | Rat | S4<br>L4-5 | R>K<br>G>S | Ref. 103 |
| TRPV1 | Human | L4-5<br>C-terminus | G>S<br>G>C<br>T>G | Ref. 104 |
| TRPV3 | Human | ANK-TM linker<br>L4-5<br>C-terminus | G>S<br>G>C<br>W>G<br>G>A<br>W>C<br>L>F | Ref. 105 |
| TRPV4, TRPV6 | Human | C-terminus | S>D<br>T>D | Mimics phosphorylation by calmodulin<br>Ref. 106 |
| TRPV4 | Human | C-terminus | Frame shift | Proposed to alter regulation by calmodulin<br>Ref. 107 |
| TRPV4 | Human | ANK | R>C<br>R>H | Ref. 108 |
| TRPY1 | Yeast | S5 | F>L | Ref. 109 |

**Table S2.**
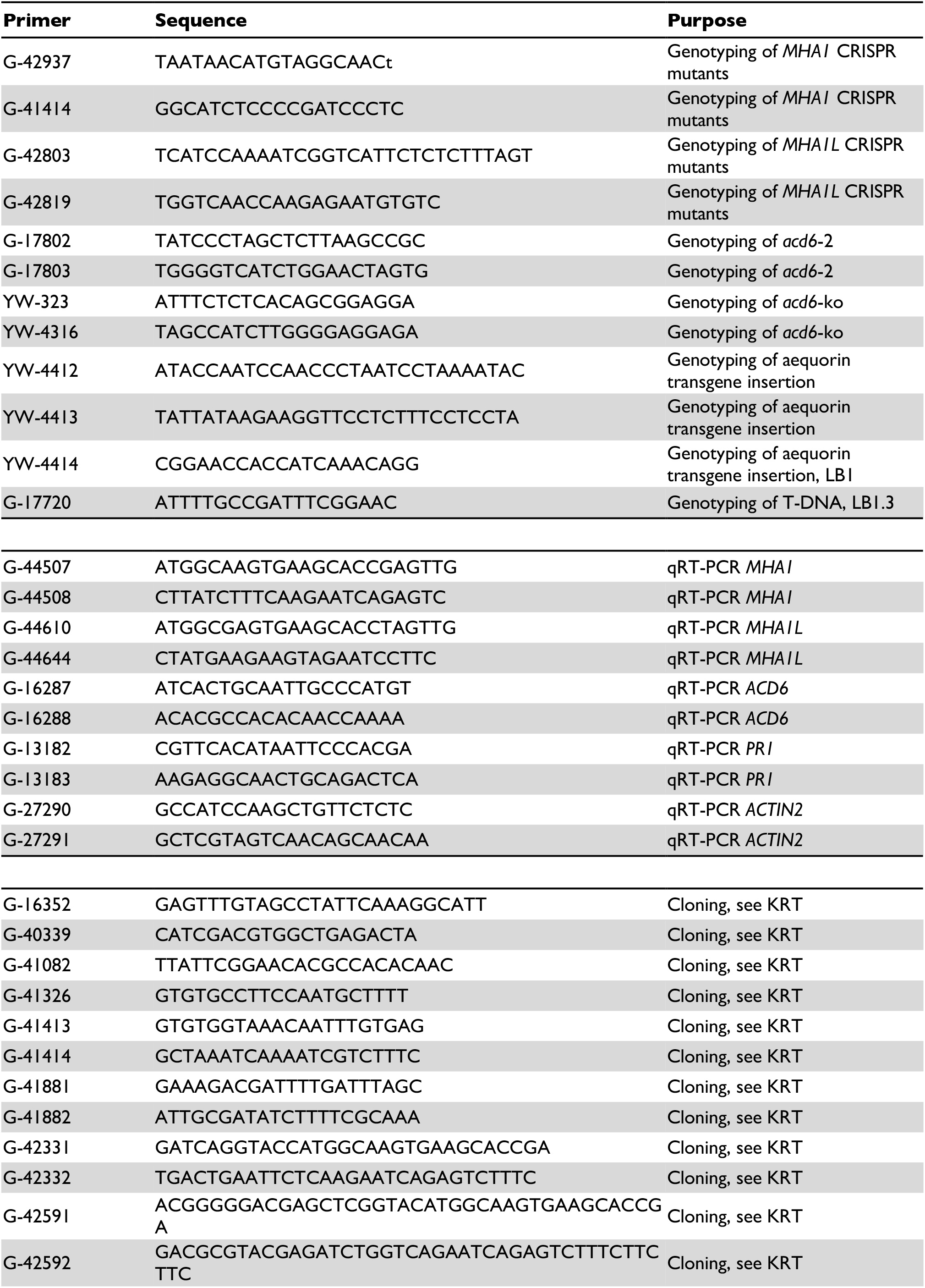
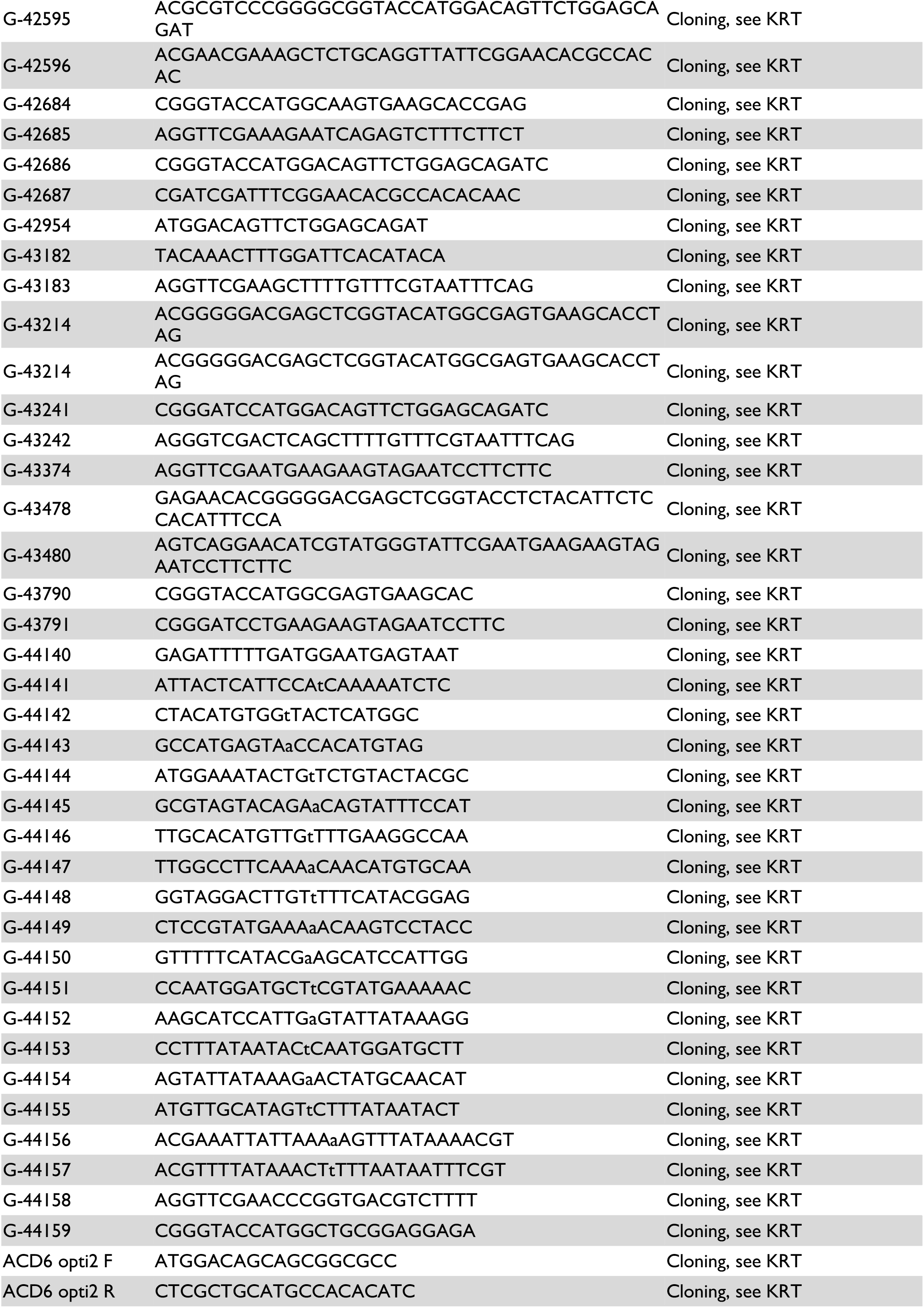
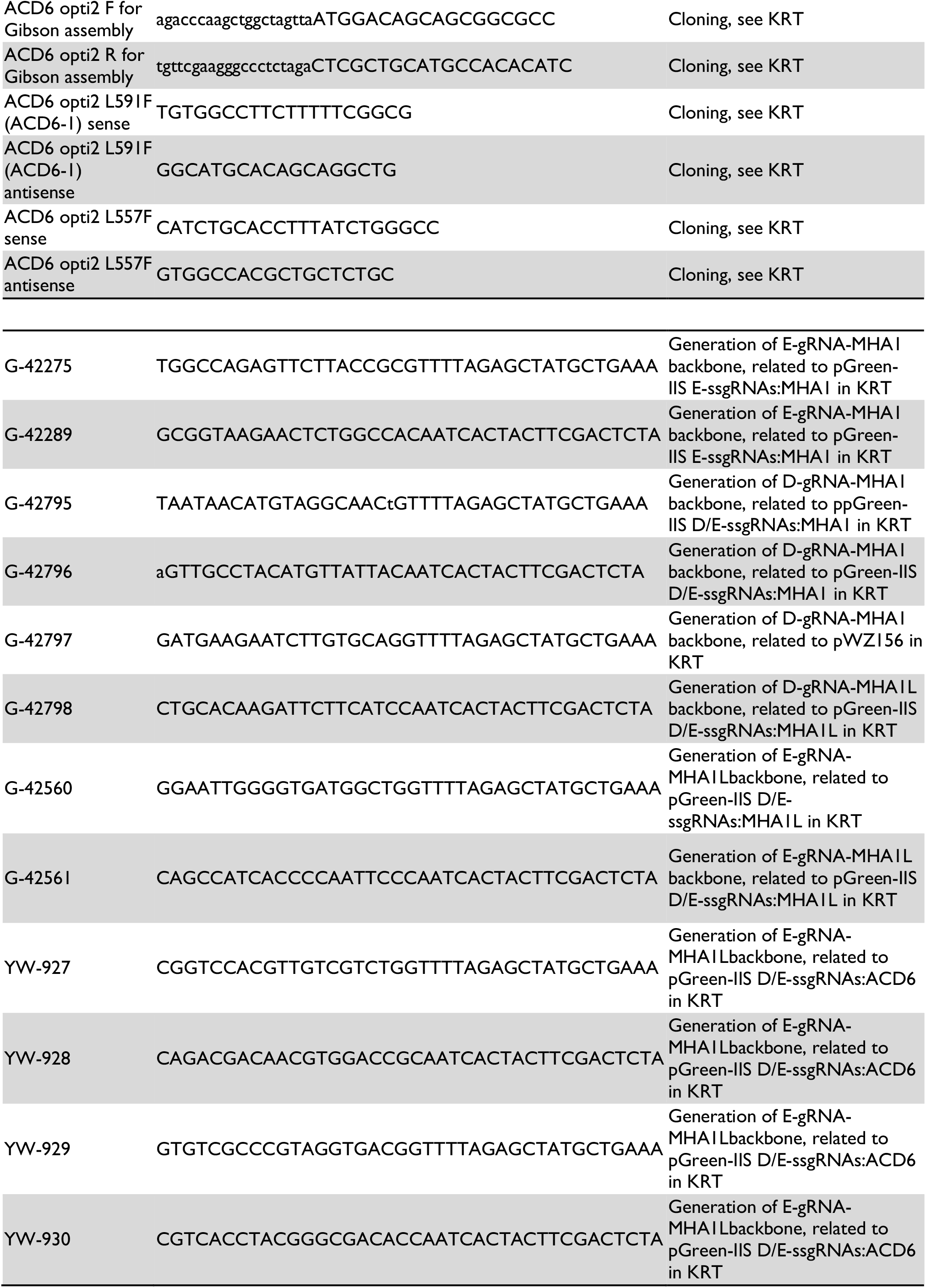
Oligonucleotide primers.

## REFERENCES

1. Todesco, M., Balasubramanian, S., Hu, T.T., Traw, M.B., Horton, M., Epple, P., Kuhns, C., Sureshkumar, S., Schwartz, C., Lanz, C., et al. (2010). Natural allelic variation underlying a major fitness trade-off in Arabidopsis thaliana. Nature 465, 632–636.

2. Todesco, M., Kim, S.-T., Chae, E., Bomblies, K., Zaidem, M., Smith, L.M., Weigel, D., and Laitinen, R.A.E. (2014). Activation of the Arabidopsis thaliana immune system by combinations of common ACD6 alleles. PLoS Genet. 10, e1004459.

3. Świadek, M., Proost, S., Sieh, D., Yu, J., Todesco, M., Jorzig, C., Rodriguez Cubillos, A.E., Plötner, B., Nikoloski, Z., Chae, E., et al. (2017). Novel allelic variants in ACD6 cause hybrid necrosis in local collection of Arabidopsis thaliana. New Phytol. 213, 900–915.

4. Rate, D.N., Cuenca, J.V., Bowman, G.R., Guttman, D.S., and Greenberg, J.T. (1999). The gain-of-function Arabidopsis acd6 mutant reveals novel regulation and function of the salicylic acid signaling pathway in controlling cell death, defenses, and cell growth. Plant Cell 11, 1695–1708.

5. Ng, G., Seabolt, S., Zhang, C., Salimian, S., Watkins, T.A., and Lu, H. (2011). Genetic dissection of salicylic acid-mediated defense signaling networks in Arabidopsis. Genetics 189, 851–859.

6. Vlot, A.C., Dempsey, D.A., and Klessig, D.F. (2009). Salicylic Acid, a multifaceted hormone to combat disease. Annu. Rev. Phytopathol. 47, 177–206.

7. Fu, Z.Q., and Dong, X. (2013). Systemic acquired resistance: turning local infection into global defense. Annu. Rev. Plant Biol. 64, 839–863.

8. Seyfferth, C., and Tsuda, K. (2014). Salicylic acid signal transduction: the initiation of biosynthesis, perception and transcriptional reprogramming. Front. Plant Sci. 5, 697.

9. Lu, H., Greenberg, J.T., and Holuigue, L. (2016). Editorial: Salicylic Acid Signaling Networks. Front. Plant Sci. 7, 238.

10. Lu, H., Rate, D.N., Song, J.T., and Greenberg, J.T. (2003). ACD6, a novel ankyrin protein, is a regulator and an effector of salicylic acid signaling in the Arabidopsis defense response. Plant Cell 15, 2408–2420.

11. Zhang, Z., Shrestha, J., Tateda, C., and Greenberg, J.T. (2014). Salicylic acid signaling controls the maturation and localization of the arabidopsis defense protein ACCELERATED CELL DEATH6. Mol. Plant 7, 1365–1383.

12. Tateda, C., Zhang, Z., Shrestha, J., Jelenska, J., Chinchilla, D., and Greenberg, J.T. (2014). Salicylic acid regulates Arabidopsis microbial pattern receptor kinase levels and signaling. Plant Cell 26, 4171–4187.

13. Yang, Y., Zhang, Y., Ding, P., Johnson, K., Li, X., and Zhang, Y. (2012). The ankyrin-repeat transmembrane protein BDA1 functions downstream of the receptor-like protein SNC2 to regulate plant immunity. Plant Physiol. 159, 1857–1865.

14. Zhang, Z., Guo, J., Zhao, Y., and Chen, J. (2019). Identification and characterization of maize ACD6-like gene reveal ZmACD6 as the maize orthologue conferring resistance to Ustilago maydis. Plant Signal. Behav. 14, e1651604.

15. Kolodziej, M.C., Singla, J., Sánchez-Martín, J., Zbinden, H., Šimková, H., Karafiátová, M., Doležel, J., Gronnier, J., Poretti, M., Glauser, G., et al. (2021). A membrane-bound ankyrin repeat protein confers race-specific leaf rust disease resistance in wheat. Nat. Commun. 12, 956.

16. Zhang, Z., Tateda, C., Jiang, S.-C., Shrestha, J., Jelenska, J., Speed, D.J., and Greenberg, J. (2017). A suite of receptor-like kinases and a putative mechano-sensitive channel are involved in autoimmunity and plasma membrane-based defenses in Arabidopsis. Mol. Plant. Microbe. Interact. 10.1094/MPMI-09-16-0184-R.

17. Zhu, W., Zaidem, M., Van de Weyer, A.-L., Gutaker, R.M., Chae, E., Kim, S.-T., Bemm, F., Li, L., Todesco, M., Schwab, R., et al. (2018). Modulation of ACD6 dependent hyperimmunity by natural alleles of an Arabidopsis thaliana NLR resistance gene. PLoS Genet. 14, e1007628.

18. Xu, G., Moeder, W., Yoshioka, K., and Shan, L. (2022). A tale of many families: calcium channels in plant immunity. Plant Cell 34, 1551–1567.

19. Bjornson, M., Pimprikar, P., Nürnberger, T., and Zipfel, C. (2021). The transcriptional landscape of Arabidopsis thaliana pattern-triggered immunity. Nat Plants 7, 579–586.

20. Tian, W., Hou, C., Ren, Z., Wang, C., Zhao, F., Dahlbeck, D., Hu, S., Zhang, L., Niu, Q., Li, L., et al. (2019). A calmodulin-gated calcium channel links pathogen patterns to plant immunity. Nature. 10.1038/s41586-019-1413-y.

21. Thor, K., Jiang, S., Michard, E., George, J., Scherzer, S., Huang, S., Dindas, J., Derbyshire, P., Leitão, N., DeFalco, T.A., et al. (2020). The calcium-permeable channel OSCA1.3 regulates plant stomatal immunity. Nature. 10.1038/s41586-020-2702-1.

22. Bi, G., Su, M., Li, N., Liang, Y., Dang, S., Xu, J., Hu, M., Wang, J., Zou, M., Deng, Y., et al. (2021). The ZAR1 resistosome is a calcium-permeable channel triggering plant immune signaling. Cell 0. 10.1016/j.cell.2021.05.003.

23. Jacob, P., Kim, N.H., Wu, F., El-Kasmi, F., Chi, Y., Walton, W.G., Furzer, O.J., Lietzan, A.D., Sunil, S., Kempthorn, K., et al. (2021). Plant “helper” immune receptors are Ca2+-permeable nonselective cation channels. Science 373, 420–425.

24. Blanc, G., Hokamp, K., and Wolfe, K.H. (2003). A recent polyploidy superimposed on older large-scale duplications in the Arabidopsis genome. Genome Res. 13, 137–144.

25. Winter, D., Vinegar, B., Nahal, H., Ammar, R., Wilson, G.V., and Provart, N.J. (2007). An “Electronic Fluorescent Pictograph” browser for exploring and analyzing large-scale biological data sets. PLoS One 2, e718.

26. Lu, H., Liu, Y., and Greenberg, J.T. (2005). Structure-function analysis of the plasma membrane-localized Arabidopsis defense component ACD6. Plant J. 44, 798–809.

27. Alcázar, R., and Parker, J.E. (2011). The impact of temperature on balancing immune responsiveness and growth in Arabidopsis. Trends Plant Sci. 16, 666–675.

28. van Wersch, R., Li, X., and Zhang, Y. (2016). Mighty Dwarfs: Arabidopsis Autoimmune Mutants and Their Usages in Genetic Dissection of Plant Immunity. Front. Plant Sci. 7, 1717.

29. Contento, A.L., and Bassham, D.C. (2012). Structure and function of endosomes in plant cells. J. Cell Sci. 125, 3511–3518.

30. Zimmermann, L., Stephens, A., Nam, S.-Z., Rau, D., Kübler, J., Lozajic, M., Gabler, F., Söding, J., Lupas, A.N., and Alva, V. (2018). A Completely Reimplemented MPI Bioinformatics Toolkit with a New HHpred Server at its Core. J. Mol. Biol. 430, 2237–2243.

31. Venkatachalam, K., and Montell, C. (2007). TRP channels. Annu. Rev. Biochem. 76, 387–417.

32. Julius, D. (2013). TRP channels and pain. Annu. Rev. Cell Dev. Biol. 29, 355–384.

33. Huffer, K.E., Aleksandrova, A.A., Jara-Oseguera, A., Forrest, L.R., and Swartz, K.J. (2020). Global alignment and assessment of TRP channel transmembrane domain structures to explore functional mechanisms. Elife 9. 10.7554/eLife.58660.

34. Becerra, C., Jahrmann, T., Puigdomènech, P., and Vicient, C.M. (2004). Ankyrin repeat-containing proteins in Arabidopsis: characterization of a novel and abundant group of genes coding ankyrin-transmembrane proteins. Gene 340, 111–121.

35. Krogh, A., Larsson, B., von Heijne, G., and Sonnhammer, E.L. (2001). Predicting transmembrane protein topology with a hidden Markov model: application to complete genomes. J. Mol. Biol. 305, 567–580.

36. Ma, W., and Berkowitz, G.A. (2007). The grateful dead: calcium and cell death in plant innate immunity. Cell. Microbiol. 9, 2571–2585.

37. Seybold, H., Trempel, F., Ranf, S., Scheel, D., Romeis, T., and Lee, J. (2014). Ca2+ signalling in plant immune response: from pattern recognition receptors to Ca2+ decoding mechanisms. New Phytol. 204, 782–790.

38. Moeder, W., Phan, V., and Yoshioka, K. (2019). Ca2+ to the rescue - Ca2+channels and signaling in plant immunity. Plant Sci. 279, 19–26.

39. Hartzell, C., Putzier, I., and Arreola, J. (2005). Calcium-activated chloride channels. Annu. Rev. Physiol. 67, 719–758.

40. Caterina, M.J., Schumacher, M.A., Tominaga, M., Rosen, T.A., Levine, J.D., and Julius, D. (1997). The capsaicin receptor: a heat-activated ion channel in the pain pathway. Nature 389, 816–824.

41. Coste, B., Mathur, J., Schmidt, M., Earley, T.J., Ranade, S., Petrus, M.J., Dubin, A.E., and Patapoutian, A. (2010). Piezo1 and Piezo2 are essential components of distinct mechanically activated cation channels. Science 330, 55–60.

42. Yuan, F., Yang, H., Xue, Y., Kong, D., Ye, R., Li, C., Zhang, J., Theprungsirikul, L., Shrift, T., Krichilsky, B., et al. (2014). OSCA1 mediates osmotic-stress-evoked Ca2+ increases vital for osmosensing in Arabidopsis. Nature 514, 367–371.

43. Kwaaitaal, M., Huisman, R., Maintz, J., Reinstädler, A., and Panstruga, R. (2011). Ionotropic glutamate receptor (iGluR)-like channels mediate MAMP-induced calcium influx in Arabidopsis thaliana. Biochem. J 440, 355–365.

44. Ranf, S., Eschen-Lippold, L., Pecher, P., Lee, J., and Scheel, D. (2011). Interplay between calcium signalling and early signalling elements during defence responses to microbe- or damage-associated molecular patterns. Plant J. 68, 100–113.

45. Choi, J., Tanaka, K., Cao, Y., Qi, Y., Qiu, J., Liang, Y., Lee, S.Y., and Stacey, G. (2014). Identification of a plant receptor for extracellular ATP. Science 343, 290–294.

46. Keinath, N.F., Waadt, R., Brugman, R., Schroeder, J.I., Grossmann, G., Schumacher, K., and Krebs, M. (2015). Live Cell Imaging with R-GECO1 Sheds Light on flg22- and Chitin-Induced Transient [Ca(2+)]cyt Patterns in Arabidopsis. Mol. Plant 8, 1188–1200.

47. Myers, B.R., Saimi, Y., Julius, D., and Kung, C. (2008). Multiple unbiased prospective screens identify TRP channels and their conserved gating elements. J. Gen. Physiol. 132, 481–486.

48. Lishko, P.V., Procko, E., Jin, X., Phelps, C.B., and Gaudet, R. (2007). The ankyrin repeats of TRPV1 bind multiple ligands and modulate channel sensitivity. Neuron 54, 905–918.

49. Qi, Z., Verma, R., Gehring, C., Yamaguchi, Y., Zhao, Y., Ryan, C.A., and Berkowitz, G.A. (2010). Ca2+ signaling by plant Arabidopsis thaliana Pep peptides depends on AtPepR1, a receptor with guanylyl cyclase activity, and cGMP-activated Ca2+ channels. Proc. Natl. Acad. Sci. U. S. A. 107, 21193–21198.

50. Bredow, M., and Monaghan, J. (2019). Regulation of Plant Immune Signaling by Calcium-Dependent Protein Kinases. Mol. Plant. Microbe. Interact. 32, 6–19.

51. Clough, S.J., Fengler, K.A., Yu, I.C., Lippok, B., Smith, R.K., Jr, and Bent, A.F. (2000). The Arabidopsis dnd1 “defense, no death” gene encodes a mutated cyclic nucleotide-gated ion channel. Proc. Natl. Acad. Sci. U. S. A. 97, 9323–9328.

52. Cheng, N.-H., Pittman, J.K., Shigaki, T., Lachmansingh, J., LeClere, S., Lahner, B., Salt, D.E., and Hirschi, K.D. (2005). Functional association of Arabidopsis CAX1 and CAX3 is required for normal growth and ion homeostasis. Plant Physiol. 138, 2048–2060.

53. Hilleary, R., Paez-Valencia, J., Vens, C., Toyota, M., Palmgren, M., and Gilroy, S. (2020). Tonoplast-localized Ca2+ pumps regulate Ca2+ signals during pattern-triggered immunity in Arabidopsis thaliana. Proc. Natl. Acad. Sci. U. S. A. 117, 18849–18857.

54. Grant, M., Brown, I., Adams, S., Knight, M., Ainslie, A., and Mansfield, J. (2000). The RPM1 plant disease resistance gene facilitates a rapid and sustained increase in cytosolic calcium that is necessary for the oxidative burst and hypersensitive cell death. Plant J. 23, 441–450.

55. Wang, J., Hu, M., Wang, J., Qi, J., Han, Z., Wang, G., Qi, Y., Wang, H.-W., Zhou, J.-M., and Chai, J. (2019). Reconstitution and structure of a plant NLR resistosome conferring immunity. Science 364, eaav5870.

56. Vo, K.T.X., Kim, C.-Y., Chandran, A.K.N., Jung, K.-H., An, G., and Jeon, J.-S. (2015). Molecular insights into the function of ankyrin proteins in plants. J. Plant Biol. 58, 271–284.

57. Knight, M.R., Campbell, A.K., Smith, S.M., and Trewavas, A.J. (1991). Transgenic plant aequorin reports the effects of touch and cold-shock and elicitors on cytoplasmic calcium. Nature.

58. Chen, H., Zou, Y., Shang, Y., Lin, H., Wang, Y., Cai, R., Tang, X., and Zhou, J.-M. (2008). Firefly luciferase complementation imaging assay for protein-protein interactions in plants. Plant Physiol. 146, 368–376.

59. Shimada, T.L., Shimada, T., and Hara-Nishimura, I. (2010). A rapid and non-destructive screenable marker, FAST, for identifying transformed seeds of Arabidopsis thaliana. Plant J. 61, 519–528.

60. Straub, T., Ludewig, U., and Neuhäuser, B. (2017). The Kinase CIPK23 Inhibits Ammonium Transport in Arabidopsis thaliana. Plant Cell 29, 409–422.

61. Grimm, D.G., Roqueiro, D., Salomé, P.A., Kleeberger, S., Greshake, B., Zhu, W., Liu, C., Lippert, C., Stegle, O., Schölkopf, B., et al. (2017). easyGWAS: A Cloud-Based Platform for Comparing the Results of Genome-Wide Association Studies. Plant Cell 29, 5–19.

62. Schindelin, J., Arganda-Carreras, I., Frise, E., Kaynig, V., Longair, M., Pietzsch, T., Preibisch, S., Rueden, C., Saalfeld, S., Schmid, B., et al. (2012). Fiji: an open-source platform for biological-image analysis. Nat. Methods 9, 676–682.

63. Waterhouse, A.M., Procter, J.B., Martin, D.M.A., Clamp, M., and Barton, G.J. (2009). Jalview Version 2--a multiple sequence alignment editor and analysis workbench. Bioinformatics 25, 1189–1191.

64. Tamura, K., Stecher, G., Peterson, D., Filipski, A., and Kumar, S. (2013). MEGA6: Molecular Evolutionary Genetics Analysis version 6.0. Mol. Biol. Evol. 30, 2725–2729.

65. Edelstein, A.D., Tsuchida, M.A., Amodaj, N., Pinkard, H., Vale, R.D., and Stuurman, N. (2014). Advanced methods of microscope control using μManager software. J Biol Methods 1. 10.14440/jbm.2014.36.

66. Schrödinger LLC (2010). The PyMOL Molecular Graphics System, Version 1.3r1.

67. Vasseur, F., Fouqueau, L., de Vienne, D., Nidelet, T., Violle, C., and Weigel, D. (2019). Nonlinear phenotypic variation uncovers the emergence of heterosis in Arabidopsis thaliana. PLoS Biol. 17, e3000214.

68. R Core Team (2019). R: A Language and Environment for Statistical Computing.

69. 1001 Genomes Consortium (2016). 1,135 Genomes Reveal the Global Pattern of Polymorphism in Arabidopsis thaliana. Cell 166, 481–491.

70. Danecek, P., Auton, A., Abecasis, G., Albers, C.A., Banks, E., DePristo, M.A., Handsaker, R.E., Lunter, G., Marth, G.T., Sherry, S.T., et al. (2011). The variant call format and VCFtools. Bioinformatics 27, 2156–2158.

71. Li, J.-F., Norville, J.E., Aach, J., McCormack, M., Zhang, D., Bush, J., Church, G.M., and Sheen, J. (2013). Multiplex and homologous recombination-mediated genome editing in Arabidopsis and Nicotiana benthamiana using guide RNA and Cas9. Nat. Biotechnol. 31, 688–691.

72. Lampropoulos, A., Sutikovic, Z., Wenzl, C., Maegele, I., Lohmann, J.U., and Forner, J. (2013). GreenGate---a novel, versatile, and efficient cloning system for plant transgenesis. PLoS One 8, e83043.

73. Gao, X., Chen, J., Dai, X., Zhang, D., and Zhao, Y. (2016). An Effective Strategy for Reliably Isolating Heritable and Cas9-Free Arabidopsis Mutants Generated by CRISPR/Cas9-Mediated Genome Editing. Plant Physiol. 171, 1794–1800.

74. Hellens, R.P., Edwards, E.A., Leyland, N.R., Bean, S., and Mullineaux, P.M. (2000). pGreen: a versatile and flexible binary Ti vector for Agrobacterium-mediated plant transformation. Plant Mol. Biol. 42, 819–832.

75. Picelli, S., Björklund, A.K., Reinius, B., Sagasser, S., Winberg, G., and Sandberg, R. (2014). Tn5 transposase and tagmentation procedures for massively scaled sequencing projects. Genome Res. 24, 2033–2040.

76. Rowan, B.A., Seymour, D.K., Chae, E., Lundberg, D.S., and Weigel, D. (2017). Methods for Genotyping-by-Sequencing. Methods Mol. Biol. 1492, 221–242.

77. Li, H., and Durbin, R. (2009). Fast and accurate short read alignment with Burrows-Wheeler transform. Bioinformatics 25, 1754–1760.

78. Li, H., Handsaker, B., Wysoker, A., Fennell, T., Ruan, J., Homer, N., Marth, G., Abecasis, G., Durbin, R., and 1000 Genome Project Data Processing Subgroup (2009). The Sequence Alignment/Map format and SAMtools. Bioinformatics 25, 2078–2079.

79. Cingolani, P., Platts, A., Wang, L.L., Coon, M., Nguyen, T., Wang, L., Land, S.J., Lu, X., and Ruden, D.M. (2012). A program for annotating and predicting the effects of single nucleotide polymorphisms, SnpEff: SNPs in the genome of Drosophila melanogaster strain w1118; iso-2; iso-3. Fly 6, 80–92.

80. Ishiga, Y., Ishiga, T., Uppalapati, S.R., and Mysore, K.S. (2011). Arabidopsis seedling flood-inoculation technique: a rapid and reliable assay for studying plant-bacterial interactions. Plant Methods 7, 32.

81. El Kasmi, F., Chung, E.-H., Anderson, R.G., Li, J., Wan, L., Eitas, T.K., Gao, Z., and Dangl, J.L. (2017). Signaling from the plasma-membrane localized plant immune receptor RPM1 requires self-association of the full-length protein. Proc. Natl. Acad. Sci. U. S. A. 114, E7385–E7394.

82. Sali, A., and Blundell, T.L. (1993). Comparative protein modelling by satisfaction of spatial restraints. J. Mol. Biol. 234, 779–815.

83. Behera, S., Zhaolong, X., Luoni, L., Bonza, M.C., Doccula, F.G., De Michelis, M.I., Morris, R.J., Schwarzländer, M., and Costa, A. (2018). Cellular Ca2+ Signals Generate Defined pH Signatures in Plants. Plant Cell 30, 2704–2719.

84. Knight, H., Trewavas, A.J., and Knight, M.R. (1996). Cold calcium signaling in Arabidopsis involves two cellular pools and a change in calcium signature after acclimation. Plant Cell 8, 489–503.

85. Mehlmer, N., Parvin, N., Hurst, C.H., Knight, M.R., Teige, M., and Vothknecht, U.C. (2012). A toolset of aequorin expression vectors for in planta studies of subcellular calcium concentrations in Arabidopsis thaliana. J. Exp. Bot. 63, 1751–1761.

86. Vang, S., Seitz, K., and Krysan, P.J. (2018). A simple microfluidic device for live cell imaging of Arabidopsis cotyledons, leaves, and seedlings. Biotechniques 64, 255–261.

87. Varadi, M., Anyango, S., Deshpande, M., Nair, S., Natassia, C., Yordanova, G., Yuan, D., Stroe, O., Wood, G., Laydon, A., et al. (2022). AlphaFold Protein Structure Database: massively expanding the structural coverage of protein-sequence space with high-accuracy models. Nucleic Acids Res. 50, D439–D444.

88. Hong, Y.S., Park, S., Geng, C., Baek, K., Bowman, J.D., Yoon, J., and Pak, W.L. (2002). Single amino acid change in the fifth transmembrane segment of the TRP Ca2+ channel causes massive degeneration of photoreceptors. J. Biol. Chem. 277, 33884–33889.

89. Kremeyer, B., Lopera, F., Cox, J.J., Momin, A., Rugiero, F., Marsh, S., Woods, C.G., Jones, N.G., Paterson, K.J., Fricker, F.R., et al. (2010). A gain-of-function mutation in TRPA1 causes familial episodic pain syndrome. Neuron 66, 671–680.

90. Van Hoeymissen, E., Held, K., Nogueira Freitas, A.C., Janssens, A., Voets, T., and Vriens, J. (2020). Gain of channel function and modified gating properties in TRPM3 mutants causing intellectual disability and epilepsy. Elife 9. 10.7554/eLife.57190.

91. Liu, H., El Zein, L., Kruse, M., Guinamard, R., Beckmann, A., Bozio, A., Kurtbay, G., Mégarbané, A., Ohmert, I., Blaysat, G., et al. (2010). Gain-of-function mutations in TRPM4 cause autosomal dominant isolated cardiac conduction disease. Circ. Cardiovasc. Genet. 3, 374–385.

92. Wang, H., Xu, Z., Lee, B.H., Vu, S., Hu, L., Lee, M., Bu, D., Cao, X., Hwang, S., Yang, Y., et al. (2019). Gain-of-Function Mutations in TRPM4 Activation Gate Cause Progressive Symmetric Erythrokeratodermia. J. Invest. Dermatol. 139, 1089–1097.

93. Jung, C., Gené, G.G., Tomás, M., Plata, C., Selent, J., Pastor, M., Fandos, C., Senti, M., Lucas, G., Elosua, R., et al. (2011). A gain-of-function SNP in TRPC4 cation channel protects against myocardial infarction. Cardiovasc. Res. 91, 465–471.

94. Winn, M.P., Conlon, P.J., Lynn, K.L., Farrington, M.K., Creazzo, T., Hawkins, A.F., Daskalakis, N., Kwan, S.Y., Ebersviller, S., Burchette, J.L., et al. (2005). A mutation in the TRPC6 cation channel causes familial focal segmental glomerulosclerosis. Science 308, 1801–1804.

95. Reiser, J., Polu, K.R., Möller, C.C., Kenlan, P., Altintas, M.M., Wei, C., Faul, C., Herbert, S., Villegas, I., Avila-Casado, C., et al. (2005). TRPC6 is a glomerular slit diaphragm-associated channel required for normal renal function. Nat. Genet. 37, 739–744.

96. Xu, H., Delling, M., Li, L., Dong, X., and Clapham, D.E. (2007). Activating mutation in a mucolipin transient receptor potential channel leads to melanocyte loss in varitint-waddler mice. Proc. Natl. Acad. Sci. U. S. A. 104, 18321–18326.

97. Kim, H.J., Li, Q., Tjon-Kon-Sang, S., So, I., Kiselyov, K., and Muallem, S. (2007). Gain-of-function mutation in TRPML3 causes the mouse Varitint-Waddler phenotype. J. Biol. Chem. 282, 36138–36142.

98. Grimm, C., Cuajungco, M.P., van Aken, A.F.J., Schnee, M., Jörs, S., Kros, C.J., Ricci, A.J., and Heller, S. (2007). A helix-breaking mutation in TRPML3 leads to constitutive activity underlying deafness in the varitint-waddler mouse. Proc. Natl. Acad. Sci. U. S. A. 104, 19583–19588.

99. Nagata, K., Zheng, L., Madathany, T., Castiglioni, A.J., Bartles, J.R., and García-Añoveros, J. (2008). The varitint-waddler (Va) deafness mutation in TRPML3 generates constitutive, inward rectifying currents and causes cell degeneration. Proc. Natl. Acad. Sci. U. S. A. 105, 353–358.

100. Dong, X.-P., Wang, X., Shen, D., Chen, S., Liu, M., Wang, Y., Mills, E., Cheng, X., Delling, M., and Xu, H. (2009). Activating mutations of the TRPML1 channel revealed by proline-scanning mutagenesis. J. Biol. Chem. 284, 32040–32052.

101. Arif Pavel, M., Lv, C., Ng, C., Yang, L., Kashyap, P., Lam, C., Valentino, V., Fung, H.Y., Campbell, T., Møller, S.G., et al. (2016). Function and regulation of TRPP2 ion channel revealed by a gain-of-function mutant. Proc. Natl. Acad. Sci. U. S. A. 113, E2363–E2372.

102. Myers, B.R., Bohlen, C.J., and Julius, D. (2008). A yeast genetic screen reveals a critical role for the pore helix domain in TRP channel gating. Neuron 58, 362–373.

103. Boukalova, S., Marsakova, L., Teisinger, J., and Vlachova, V. (2010). Conserved residues within the putative S4-S5 region serve distinct functions among thermosensitive vanilloid transient receptor potential (TRPV) channels. J. Biol. Chem. 285, 41455–41462.

104. Lin, Z., Chen, Q., Lee, M., Cao, X., Zhang, J., Ma, D., Chen, L., Hu, X., Wang, H., Wang, X., et al. (2012). Exome sequencing reveals mutations in TRPV3 as a cause of Olmsted syndrome. Am. J. Hum. Genet. 90, 558–564.

105. Wang, G., and Wang, K. (2017). The Ca2+-Permeable Cation Transient Receptor Potential TRPV3 Channel: An Emerging Pivotal Target for Itch and Skin Diseases. Mol. Pharmacol. 92, 193–200.

106. Arbabian, A., Iftinca, M., Altier, C., Singh, P.P., Isambert, H., and Coscoy, S. (2020). Mutations in calmodulin-binding domains of TRPV4/6 channels confer invasive properties to colon adenocarcinoma cells. Channels 14, 101–109.

107. Mah, W., Sonkusare, S.K., Wang, T., Azeddine, B., Pupavac, M., Carrot-Zhang, J., Hong, K., Majewski, J., Harvey, E.J., Russell, L., et al. (2016). Gain-of-function mutation in TRPV4 identified in patients with osteonecrosis of the femoral head. J. Med. Genet. 53, 705–709.

108. Landouré, G., Zdebik, A.A., Martinez, T.L., Burnett, B.G., Stanescu, H.C., Inada, H., Shi, Y., Taye, A.A., Kong, L., Munns, C.H., et al. (2010). Mutations in TRPV4 cause Charcot-Marie-Tooth disease type 2C. Nat. Genet. 42, 170–174.

109. Su, Z., Zhou, X., Haynes, W.J., Loukin, S.H., Anishkin, A., Saimi, Y., and Kung, C. (2007). Yeast gain-of-function mutations reveal structure-function relationships conserved among different subfamilies of transient receptor potential channels. Proc. Natl. Acad. Sci. U. S. A. 104, 19607–19612.

